# The extracellular loops of *Salmonella* Typhimurium outer membrane protein A (OmpA) maintain the stability of *Salmonella* containing vacuole (SCV) in murine macrophages and protect the bacteria from autophagy-dependent lysosomal degradation

**DOI:** 10.1101/2021.11.07.467609

**Authors:** Atish Roy Chowdhury, Dipasree Hajra, Dipshikha Chakravortty

## Abstract

After entering the host cells, *Salmonella* Typhimurium (STM) stays inside a modified membrane-bound compartment called *Salmonella* containing vacuole (SCV). The biogenesis and stability of SCV are crucial for the intracellular proliferation of *Salmonella*. Our research has provided a novel mechanistic view on the role of a bacterial porin OmpA in maintaining the stability of SCV. We found that the deletion of OmpA forces the bacteria to escape from the SCV during the immediate early stage of infection. In the absence of OmpA, the bacteria failed to retain the LAMP-1 and came into the host cell’s cytosol. Subsequently, the cytosolic population of STM *ΔompA* activated the host autophagy machinery after colocalizing with syntaxin 17 and LC3B. The autophagosomes carrying STM *ΔompA* were targeted to the lysosomes for degradation. Inhibition of autophagy pathway using bafilomycin A1 restored the intracellular proliferation of STM *ΔompA*. We further showed that the four extracellular loops of OmpA played a crucial role in holding the LAMP-1 pool around the SCV. We have altered the extracellular loop sequences of *Salmonella* OmpA by site-directed mutagenesis and observed that the bacteria failed to maintain the LAMP-1 pool around the SCV, which finally resulted in their release into the cytosol of the host macrophages. Surprisingly, the cytosolic population of *Salmonella* having mutations in the extracellular loops of OmpA didn’t activate the lysosomal degradation pathway like STM *ΔompA,* which helped them to survive within the murine macrophages. In summary, our study revealed an OmpA dependent novel strategy utilized by *Salmonella* to combat host autophagy by promoting the stability of SCV.

## Introduction

*Salmonella* Typhimurium (STM), the non-typhoidal serovar of the enteric pathogen *Salmonella enterica,* causes self-limiting diarrheal disease and gastroenteritis in humans [1]. It causes typhoid fever-like symptoms in mice and is considered an excellent model to study the pathogenesis of the human pathogen *Salmonella* Typhi. The pathogen enters the body of healthy individuals with contaminated food and water. The Global Burden of Diseases, Injuries, and Risk Factors Study (GBD) estimated an occurrence of 5,35,000 cases of invasive non-typhoidal *Salmonella* infection worldwide, with approximately 77500 deaths in 2017 [2]. *Salmonella* Typhimurium can invade a wide range of host cells. After entering the host cells, *Salmonella* resides within a modified membrane-bound acidic compartment called *Salmonella* containing vacuole (SCV) [3–5]. The successful intracellular proliferation of the bacteria depends upon the formation and maintenance of intact SCV within the host cells. The disruption of SCV imposes a dramatic outcome on the fate of the intracellular bacteria. The leakage of SCV and subsequent release of the bacteria into the cytosol of phagocytic cells abrogate bacterial proliferation. In contrast, the cytosol of the non-phagocytic epithelial cells promotes bacterial proliferation [5–9]. *Salmonella* majorly uses SPI-1 and SPI-2 encoded virulent factors to control the biogenesis of SCV. These SPI-encoded virulent effectors work in concert with host proteins to maintain the stability of SCV [4, 10–12]. However, the contributions of non-SPI virulent genes in the biogenesis and stability of SCV have been poorly understood.

Outer membrane protein A (OmpA) is a β barrel porin protein found on the outer membrane of *Salmonella* Typhimurium. It consists of eight anti-parallel β sheets, connected by four externally exposed extracellular loops and four periplasmic turns. With the help of its periplasmic domain, it protects *Salmonella* from oxidative stress by changing the outer membrane permeability [13]. The deficiency of OmpA compromises the biofilm-forming ability of the bacteria in response to bile salt stress [14]. In our previous study, we have proved that deletion of OmpA significantly hampers the stability of the outer membrane of *Salmonella*, which makes the bacteria susceptible to *in vivo* nitrosative stress[15]. We have also found that *ompA* knockout *Salmonella* quits the SCV during the late phase of infection in murine macrophages, suggesting a previously unknown role of OmpA in regulating the stability of SCV.

In this current study, we addressed a novel role of *Salmonella* Typhimurium OmpA in maintaining SCV stability. We have shown that the cytosolic population of *Salmonella* lacking OmpA activates the host autophagy machinery and is cleared by the lysosomal degradative pathway. Our study further revealed a strong interaction between the OmpA of intracellular *Salmonella* with host LAMP-1 in macrophages. By introducing mutations in the externally exposed extracellular loops of OmpA, we dissected the role of *Salmonella* OmpA in modulating the intracellular vacuolar life of the pathogen. To the best of our knowledge, this is the first study illustrating the precise role of the extracellular loops of *Salmonella* Typhimurium OmpA in the intracellular virulence of bacteria.

## Results

### OmpA deficient strain of *Salmonella* quits SCV during the late phase of infection in murine macrophages and activates host autophagy machinery

Intracellular *Salmonella* Typhimurium residing within the SCV can inhibit phagolysosome maturation [4]. The successful systemic colonization of *Salmonella* Typhimurium depends upon forming a replicative niche within the host cells [16]. However, the biogenesis of SCV is a complicated phenomenon. *Salmonella* employs a plethora of proteins and virulent factors that work in conjunction with host factors for the construction of SCV. Most of the studies that have addressed the intracellular pathogenesis of *Salmonella* have discovered the role of SPI-1 and SPI-2 encoded type 3 secretion systems (T3SS1 and T3SS2) and virulent factors in SCV biogenesis [17]. Earlier, our group has reported the role of outer membrane protein A (OmpA) in maintaining the stability of SCV within murine macrophages[15]. To validate our previous observation, the vacuolar niche of the wild type, *ompA* deficient, and complemented strains of *S*. Typhimurium was checked in murine macrophages. STM *ΔompA* was found to be residing with in the cytoplasm of RAW264.7 cells during the late phase of infection (**Figure 1A**). The poor colocalization of STM *ΔompA* with LAMP-1 proved their release from the SCV (**Figure 4.1B**). When the *ompA* gene was complemented in the knockout bacteria, there was a reversal in the vacuolar escaping phenotype (**Figure 1A and 1B**). Wild type *Salmonella* can recruit a non-receptor tyrosine kinase named focal adhesion kinase (FAK) on the surface of the SCV in an SPI-2 encoded T3SS2 dependent manner. FAK can suppress the host autophagy machinery by activating the Akt-mTOR signaling pathway [18, 19]. Our previous study proved that intracellular STM *ΔompA* is unable to produce and secrete the SPI-2 encoded translocon proteins into the host cell’s cytosol [15], suggesting the formation of a malfunctioning T3SS2. Hence, we have hypothesized that the infection of macrophages with STM *ΔompA* may activate host autophagy machinery. RAW264.7 cells were infected with STM (WT) and *ΔompA* to evaluate the recruitment of autophagy markers (syntaxin 17 and LC3B) around the bacteria during the late phase of infection. Syntaxin 17 is an autophagosomal SNARE protein with a unique C-terminal hairpin structure of two tandem trans-membrane domains which are constructed with glycine zipper motifs and interacts with the autophagosomal membrane [20]. Syntaxin 17 can further recruit SNAP 29 and lysosomal SNARE protein VAMP8 [21]. It helps in the fusion of the autophagosome with lysosome and degradation of enclosed contents [22]. Microtubule-associated protein 1A/ 1B-light chain 3 (MAP-LC3/ LC3/ Atg8), which loads the cargo in the autophagosome, is considered one of the important autophagy markers. Usually, LC3B is diffused throughout the cytosol. Upon autophagy initiation, LC3B is cleaved by cysteine protease Atg4 to form LC3B-I and further modified by the association of phosphatidylethanolamine to create LC3B-II. This lipidated form of LC3B (LC3B-II) forms distinct small puncta in the cytosol and helps to seal the membrane of autophagosome carrying cargo [23]. In line with our expectation, an increased recruitment of syntaxin 17 (**Figure 1C and 1D**) and LC3B (**Figure 1E and 1F**) was observed around STM *ΔompA* compared to the wild type bacteria during the late phase of infection. This suggests that STM *ΔompA* can damage SCV and eventually activates host autophagy machinery in macrophages. To firmly support this conclusion, the co-staining of LAMP-1 and syntaxin 17 was performed in RAW264.7 cells infected with STM (WT) and *ΔompA* (**Figure S1A and S1B**). It was found that the wild type *Salmonella* staying within intact SCV (**Figure S1A.1**) can restrict the recruitment of syntaxin 17 (**Figure S1A.3 and S1A.5**). In the contrary, the majority of intracellular STM *ΔompA* that hardly colocalize with LAMP-1 (**Figure S1A.2**) profoundly sequester syntaxin 17 (**Figure S1A.4 and S1A.6**). The better colocalization of autophagy markers with STM *ΔompA* (**Figure 1D and 1F**) suggested the formation of syntaxin 17^+^ LC3B^+^ autophagosome around the mutant bacteria and subsequent activation of host autophagy machinery (xenophagy). Autophagy can target the pathogen trapped inside the autophagosome to the lysosomes for degradation. We verified this hypothesis by measuring the activation and subsequent fusion of the lysosomes with wild type and *ompA* deficient *Salmonella.* Before infecting the cells, the lysosomes were loaded with Texas red ovalbumin, and their colocalization with the intracellular pathogens was evaluated. Compared to STM (WT), the enhanced colocalization of STM *ΔompA* with Texas red (trapped inside lysosomes) suggested a sharp rise in lysosomal activity (**Figure 1G and 1H**). As a control, STM (WT): *LLO*, a wild type bacterial strain that leaves SCV because of the expression of pore-forming toxin listeriolysin O from *Listeria monocytogenes,* was used. The reduced colocalization of STM (WT): *LLO* with lysosomes robustly proves the role of OmpA to prevent lysosomal fusion with the cytosolic pool of wild type bacteria (**Figure 1G and 1H**). To prove the subsequent activation of autophagy and lysosomal degradation upon infection of macrophages with STM *ΔompA*, the colocalization between host syntaxin 17 and Texas red ovalbumin was studied. It was found that unlike the wild type *Salmonella* (**Figure S2A.1, S2A.3, and S2A.5**), STM *ΔompA* simultaneously colocalizes with both syntaxin 17 (**Figure S2A.2**) and lysosomes (**Figure S2A.4 and S2A.6**). This suggests that the syntaxin 17^+^ autophagosome carrying STM *ΔompA* is finally targeted to the lysosome for degradation. The enhanced activity of lysosomes upon ingestion of any cargo can be estimated by measuring the activity of lysosomal enzymes such as acid phosphatases [24, 25]. The intense activity of lysosomal enzymes upon infection of macrophages with STM *ΔompA* was measured by acid phosphatase assay. The lysosomal acid phosphatase activity was found impaired when the cells were infected with the wild type and complemented strains of *Salmonella* (**Figure 1I**). The improved activity of acid phosphatases from the cells infected with STM *ΔompA* (**Figure 1I**) suggested an active function of lysosomes in killing the pathogen. Taken together, our data proves that during the late phase of infection in murine macrophages, STM *ΔompA* reaches the cytosol of the host cell and activates host autophagy machinery. As a result, the mutant bacteria trapped inside syntaxin 17^+^, LC3B^+^ autophagosome is targeted to the lysosomal degradation pathway, which might be a reason behind the clearance of the bacteria from macrophages.

**Figure 1.**
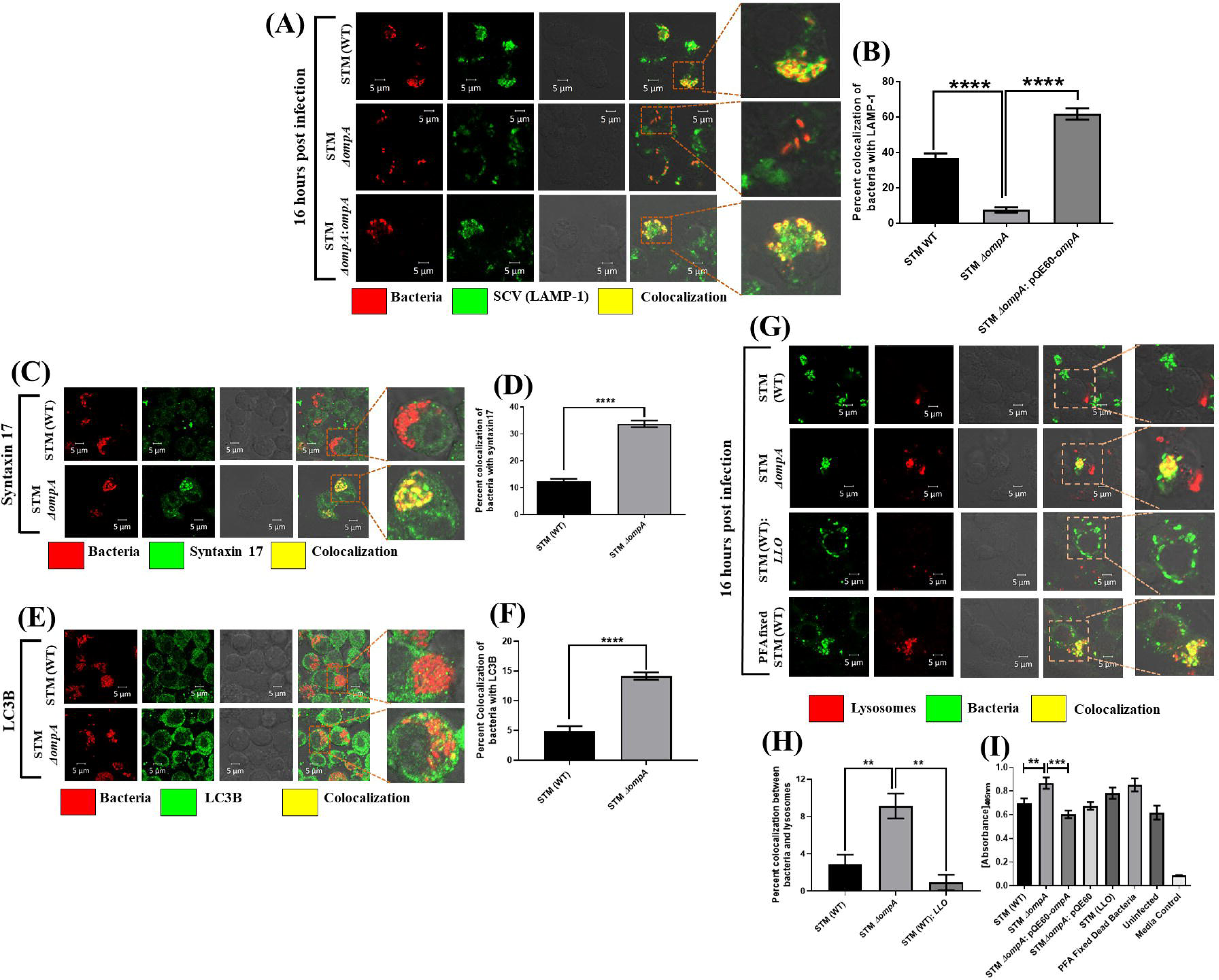
OmpA deficient strain of *Salmonella* quits SCV during the late phase of infection in murine macrophages and activates host autophagy machinery. (A) RAW264.7 cells were infected with STM (WT): RFP, *ΔompA*: RFP, and *ΔompA*: pQE60- *ompA* at MOI of 20. Cells were fixed at 16 hours post-infection & LAMP-1 was labeled with anti-mouse LAMP-1 antibody. To stain the complemented strain anti-*Salmonella* antibody was used. (B) The quantification of LAMP-1 recruitment on bacteria in RAW 264.7 cells has been represented in a graph. Percent colocalization was determined after analyzing more than 60 different microscopic stacks from two independent experiments. Scale bar = 5μm, [n≥60, N=2]. (C-F) RAW264.7 cells were infected with STM (WT): *RFP* and *ΔompA*: *RFP* at MOI of 20. Cells were fixed at 16 hours post-infection. Two autophagy markers, (C) syntaxin 17 and (E) LC3B, were stained with rabbit-raised anti-mouse syntaxin 17 and LC3B-II primary antibodies, respectively. The quantification of (D) syntaxin 17 and (F) LC3B recruitment on STM (WT): *RFP* and *ΔompA*: *RFP* has been represented in the form of two graphs. (D) The percent colocalization of syntaxin 17 with bacteria was determined after analyzing 100 microscopic stacks from three independent experiments (n=100. N=3). (F) The percent colocalization between the bacteria and LC3B was determined after analyzing more than 60 different microscopic stacks from two independent experiments. Scale bar = 5μm, [n≥60, N=2]. To stain the lysosomes, RAW 264.7 cells were pre-treated with Texas red ovalbumin for 30 minutes. (G) The cells are washed thrice with PBS after that and infected with STM (WT): *GFP*, *ΔompA*: *GFP*, and STM (WT): *LLO*, respectively, at MOI of 20. PFA-fixed dead bacteria were used for infection at MOI of 25. To stain STM (WT): *LLO* and PFA fixed dead bacteria rabbit-anti *Salmonella* O primary and anti-rabbit dylight 488 secondary antibodies were used. (H) The colocalization of lysosomes with bacteria has been represented in the form of a graph. The percent colocalization between Texas red and the bacteria was determined after counting 50 microscopic stacks from two independent experiments (n=50, N=2). Scale bar = 5μm. (I) To measure the acid phosphatase activity of lysosomes, RAW 264.7 cells were infected with STM (WT), *ΔompA*, *ΔompA*: pQE60-*ompA*, *ΔompA*: pQE60, STM (WT): *LLO*, and PFA fixed dead bacteria at MOI of 10. Twelve hours post-infection, the cells were washed with PBS and further incubated for 4 hours at 37^0^C with a buffer containing sodium acetate, triton-X-100, and *p-*nitrophenyl phosphate (pNPP). The absorbance of the supernatant was measured at 405 nm using a microplate reader (n=6, N=2). ***(p) **< 0.005*, *(p) ***< 0.0005*, *(p) ****< 0.0001* (Student’s *t*-test).**

**Figure 2.**
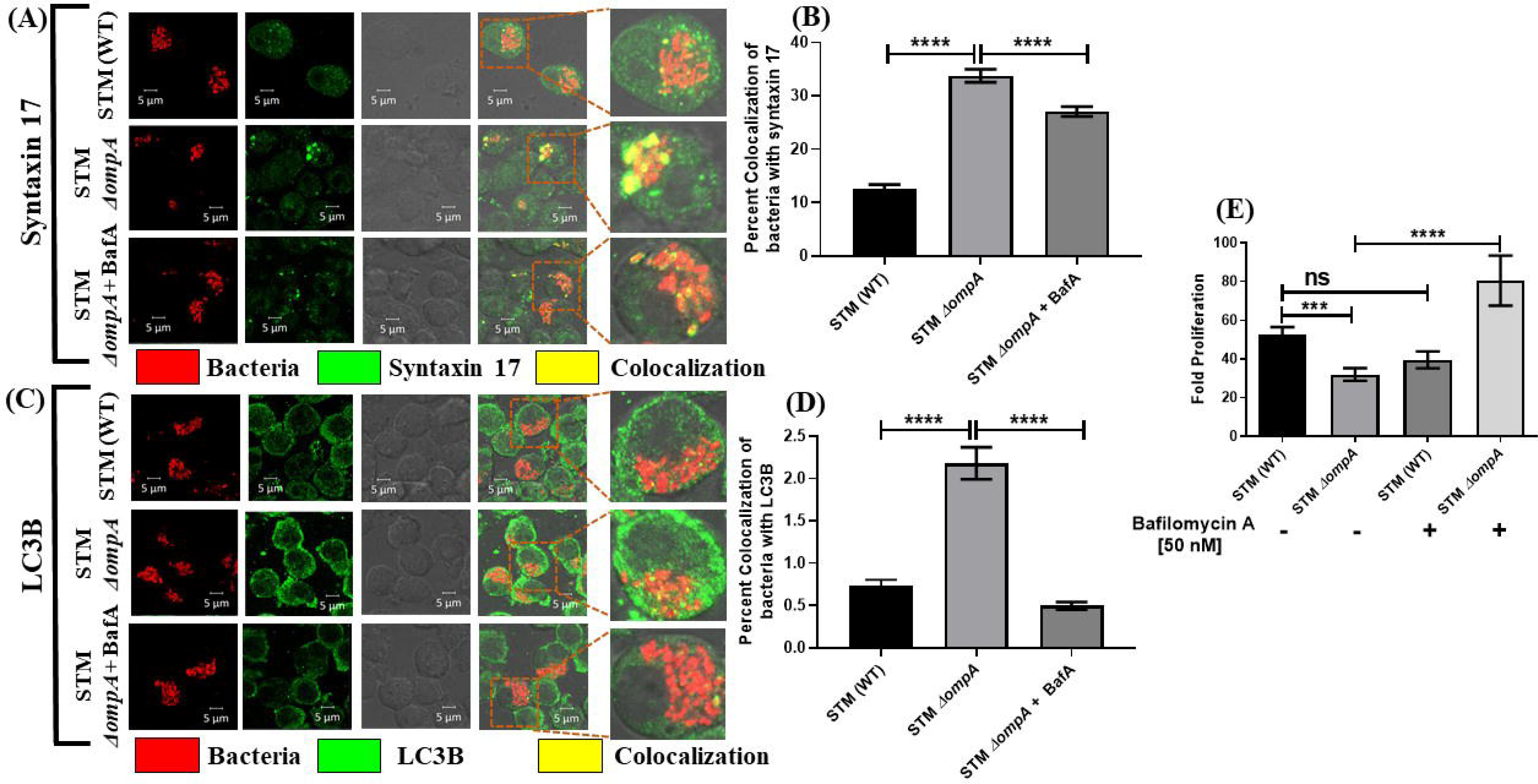
Inhibition of host autophagy using bafilomycin A restored the intracellular proliferation of *ompA* deficient strain of *Salmonella* Typhimurium. RAW264.7 cells were infected with STM (WT): *RFP* and *ΔompA*: *RFP* at MOI of 20. One set of cells infected with STM *ΔompA*: *RFP* were treated with bafilomycin A (50 nM). Cells were fixed at 16 hours post-infection. Two autophagy markers, (A) syntaxin 17 and (C) LC3B, were stained with rabbit-raised anti-mouse syntaxin 17 and LC3B-II primary antibodies, respectively. The quantification of (B) syntaxin 17 and (D) LC3B recruitment on STM (WT): *RFP*, *ΔompA*: *RFP*, and *ΔompA*: *RFP* under bafilomycin A treatment have been represented in the form of two graphs. (B) The percent colocalization of syntaxin 17 with bacteria was determined after analyzing 100 microscopic stacks from three independent experiments (n=100. N=3). (D) The percent colocalization between the bacteria and LC3B was determined after analyzing more than 60 different microscopic stacks from two independent experiments. Scale bar = 5μm, [n≥60, N=2]. (E) Intracellular survival of STM (WT) and *ΔompA* (MOI-10) in RAW264.7 cells (16 hours post-infection) in presence and absence of autophagy inhibitor bafilomycin A (50 nM). The bacteria’s fold proliferation was calculated by normalizing the CFU at 16 hours to CFU at 2 hours (n=3, N=2). ***(p) ***< 0.0005*, *(p) ****< 0.0001*, ns= non-significant (Student’s *t*-test).**

**Figure 3.**
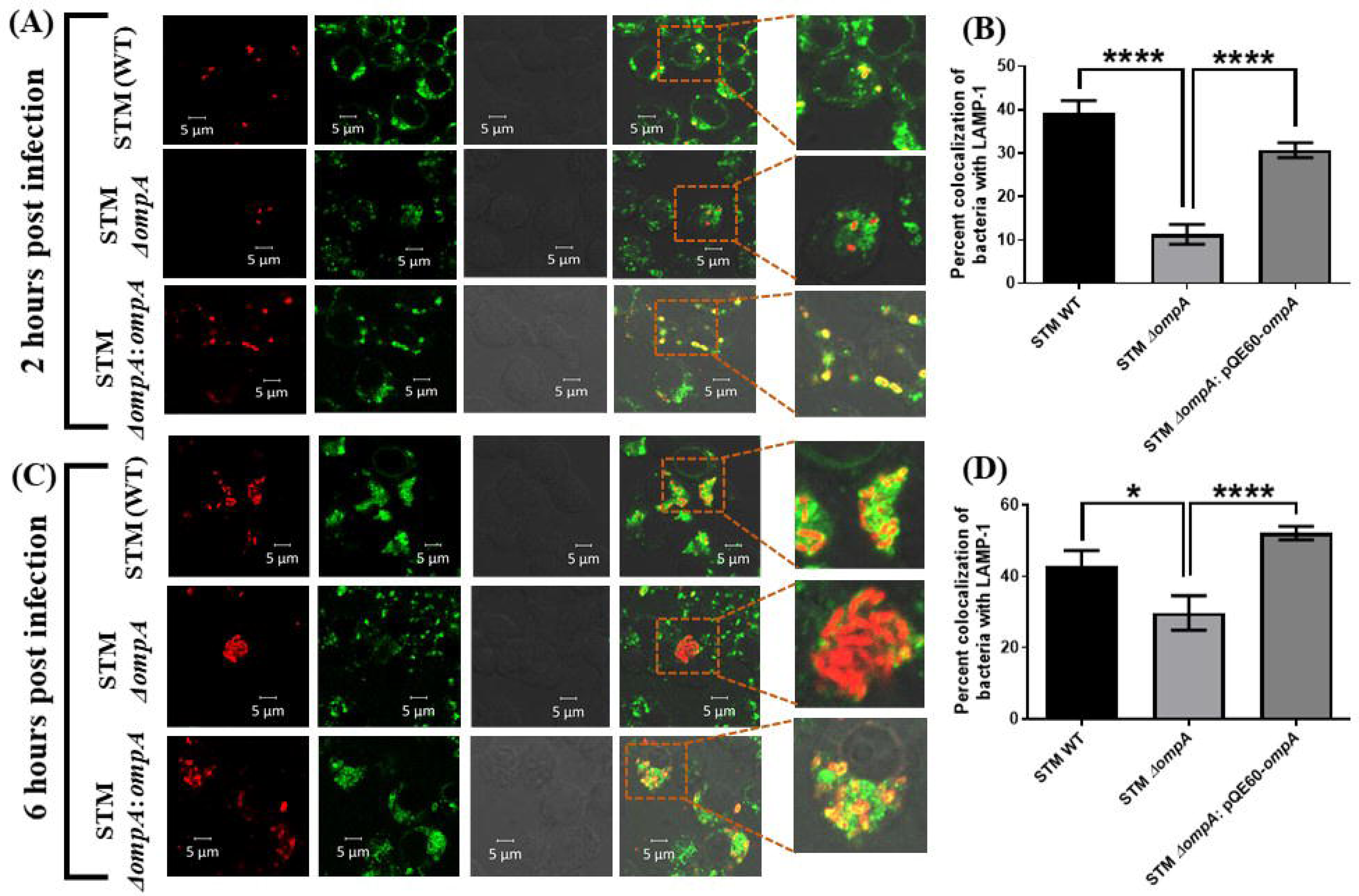
STM *ΔompA* quits the SCV in murine macrophages before the early stage of infection. (A-D) RAW264.7 cells were infected with STM (WT): RFP, *ΔompA*: RFP, and *ΔompA*: pQE60-*ompA* at MOI of 20. Cells were fixed at (A) 2 hours (early phase) and (C) 6 hours (middle phase) post-infection & LAMP-1 was labeled with anti-mouse LAMP-1 antibody. To stain the complemented strain and the PFA fixed dead bacteria anti-*Salmonella* antibody was used. The quantification of LAMP-1 recruitment on bacteria in RAW 264.7 cells at (B) 2 hours and (D) 6 hours post-infection has been represented in the form of two graphs. (B) During the early stage of infection (2 hours post-infection), the percent colocalization of bacteria with LAMP-1 was determined after analyzing more than 50 different microscopic stacks from two independent experiments [n≥50, N=2]. (D) During the middle stage of infection (6 hours post-infection), the percent colocalization of bacteria with LAMP-1 was determined after analyzing more than 40 different microscopic fields from two independent experiments [n≥40, N=2]. Scale bar = 5μm. ***(P)* *< 0.05, *(P)* ****< 0.0001, ns= non-significant, (Student’s *t-*test).**

**Figure 4.**
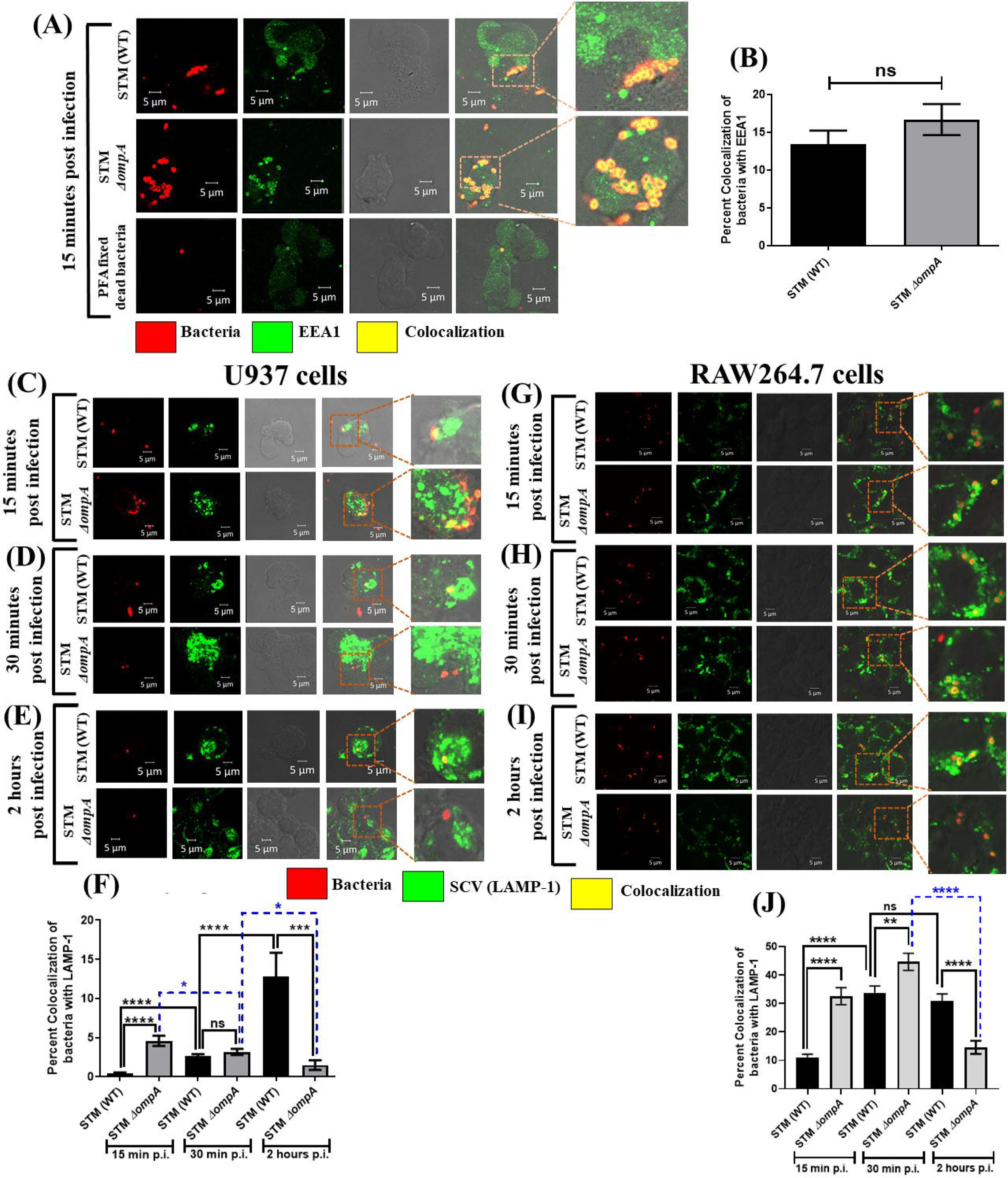
STM *ΔompA* quits the SCV during the immediate early stage of infection in macrophages. PMA activated U937 cells were infected with STM (WT), *ΔompA*, and PFA fixed dead bacteria at MOI of 25. (A) Cells were fixed at 15 minutes post-infection (immediate early stage of infection). The bacteria have been stained with rabbit-raised anti-*Salmonella* O primary antibody. Early endosome antigen (EEA1) was labeled with an anti-human EEA1 antibody raised in the mouse. (B) The quantification of EEA1 recruitment on bacteria in U937 cells has been represented in the form of three graphs. During the immediate early stage of infection (15 minutes post-infection), the percent colocalization of bacteria with EEA1 was determined after analyzing more than 30 different microscopic stacks from two independent experiments [n≥30, N=2]. PMA activated U937 cells were infected with STM (WT): RFP, and *ΔompA*: RFP at MOI of 20. Cells were fixed at (C) 15 minutes (immediate early phase), (D) 30 minutes, and (E) 2 hours post-infection & LAMP-1 was labeled with anti-human LAMP-1 antibody. (F) The quantification of LAMP-1 recruitment on bacteria in U937 cells at 15 minutes, 30 minutes, and 2 hours post-infection has been represented in the form of a graph. (F) The percent colocalization of bacteria with LAMP-1 was determined after analyzing more than 50 different microscopic stacks from two independent experiments [n≥50, N=2]. Scale bar = 5μm. RAW264.7 cells were infected with STM (WT): RFP, and *ΔompA*: RFP at MOI of 20. Cells were fixed at (G) 15 minutes (immediate early phase), (H) 30 minutes, and (I) 2 hours post-infection & LAMP-1 was labeled with anti-mouse LAMP-1 antibody. (J) The quantification of LAMP-1 recruitment on bacteria in RAW264.7 cells at 15 minutes, 30 minutes, and 2 hours post-infection has been represented in the form of a graph. (J) The percent colocalization of bacteria with LAMP-1 was determined after analyzing more than 60 different microscopic stacks from two independent experiments [n≥60, N=2]. Scale bar = 5μm. ***(P)* **< 0.005, *(P)* ***< 0.0005, *(P)* ****< 0.0001, ns= non-significant, (Student’s *t-*test).**

**Figure 5.**
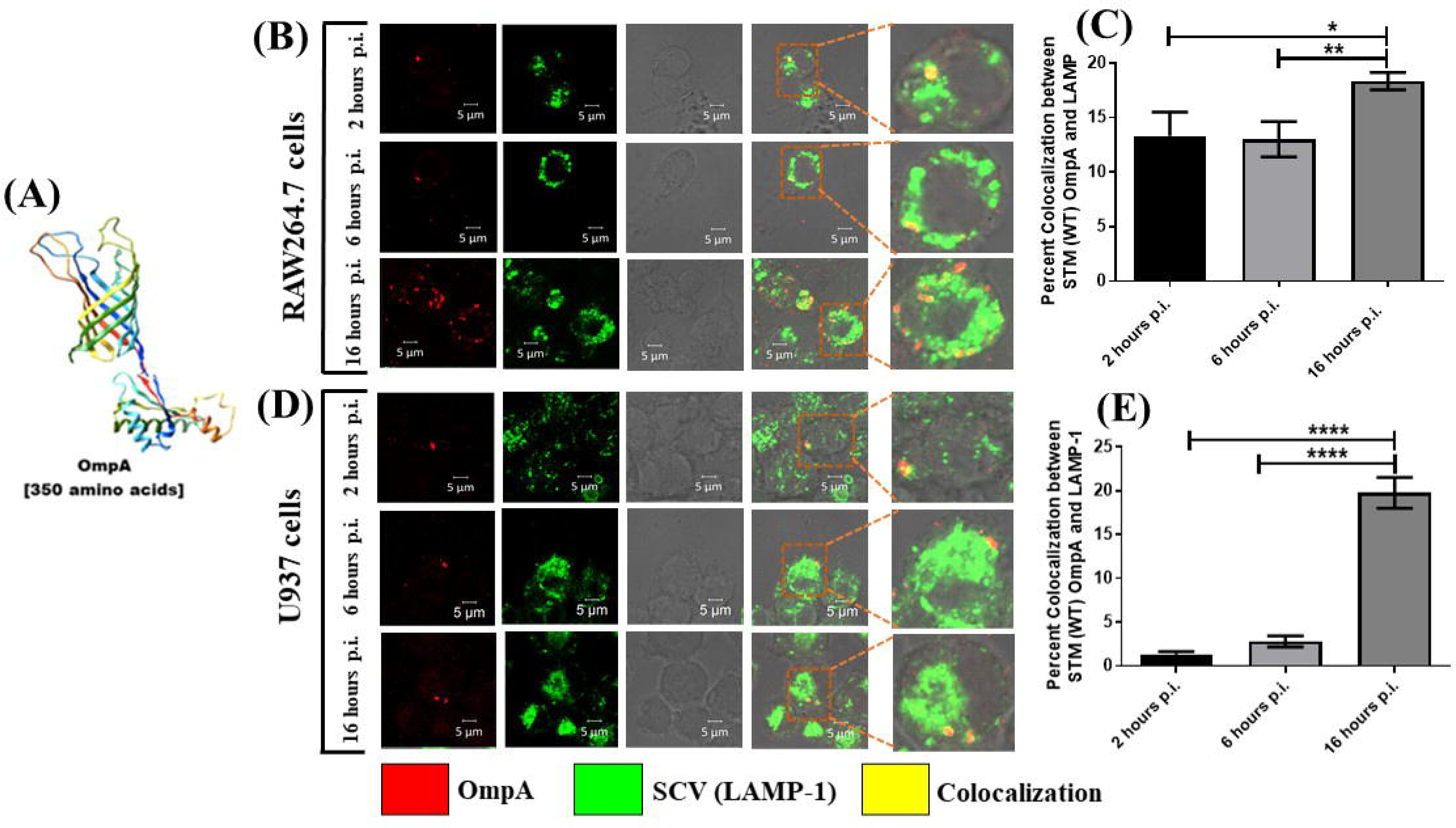
*Salmonella* Typhimurium OmpA plays a critical role in maintaining the interaction with host LAMP-1 (SCV) during infection in macrophages. (A) The structure of *Salmonella* Typhimurium OmpA obtained with the help of SWISS-MODEL software (B) RAW 264.7 cells were infected with STM (WT) at MOI of 20. Cells were fixed at 2 hours (early phase), 6 hours (middle phase), and 16 hours (late phase) post-infection. Intracellular *Salmonella* was stained with rabbit-raised anti-*Salmonella* OmpA primary antibody. In RAW264.7 cells, LAMP-1 was labeled with rat-raised anti-mouse LAMP-1 antibody. (C) The quantification of LAMP-1 recruitment on bacteria in RAW264.7 cells at 2 hours, 6 hours, and 16 hours post-infection has been represented in the form of a graph. (C) The percent colocalization of bacteria with LAMP-1 was determined after analyzing 50 different microscopic stacks from two independent experiments [n=50, N=2]. (D) PMA activated U937 cells were infected with STM (WT) at an MOI of 20. Cells were fixed at 2 hours (early phase), 6 hours (middle phase), and 16 hours (late phase) post-infection. Intracellular *Salmonella* was stained with rabbit-raised anti-*Salmonella* OmpA primary antibody. In U937 cells, LAMP-1 was labeled with an anti-human LAMP-1 antibody. (E) The quantification of LAMP-1 recruitment on bacteria in U937 cells at 2 hours, 6 hours, and 16 hours post-infection has been represented in the form of a graph. (E) The percent colocalization of bacteria with LAMP-1 was determined after analyzing 50 different microscopic stacks from three independent experiments [n=50, N=3]. Scale bar = 5μm. ***(P)* *< 0.05, *(P)* **< 0.005, *(P)* ****< 0.0001, ns= non-significant, (Student’s *t-*test).**

**Figure 6.**
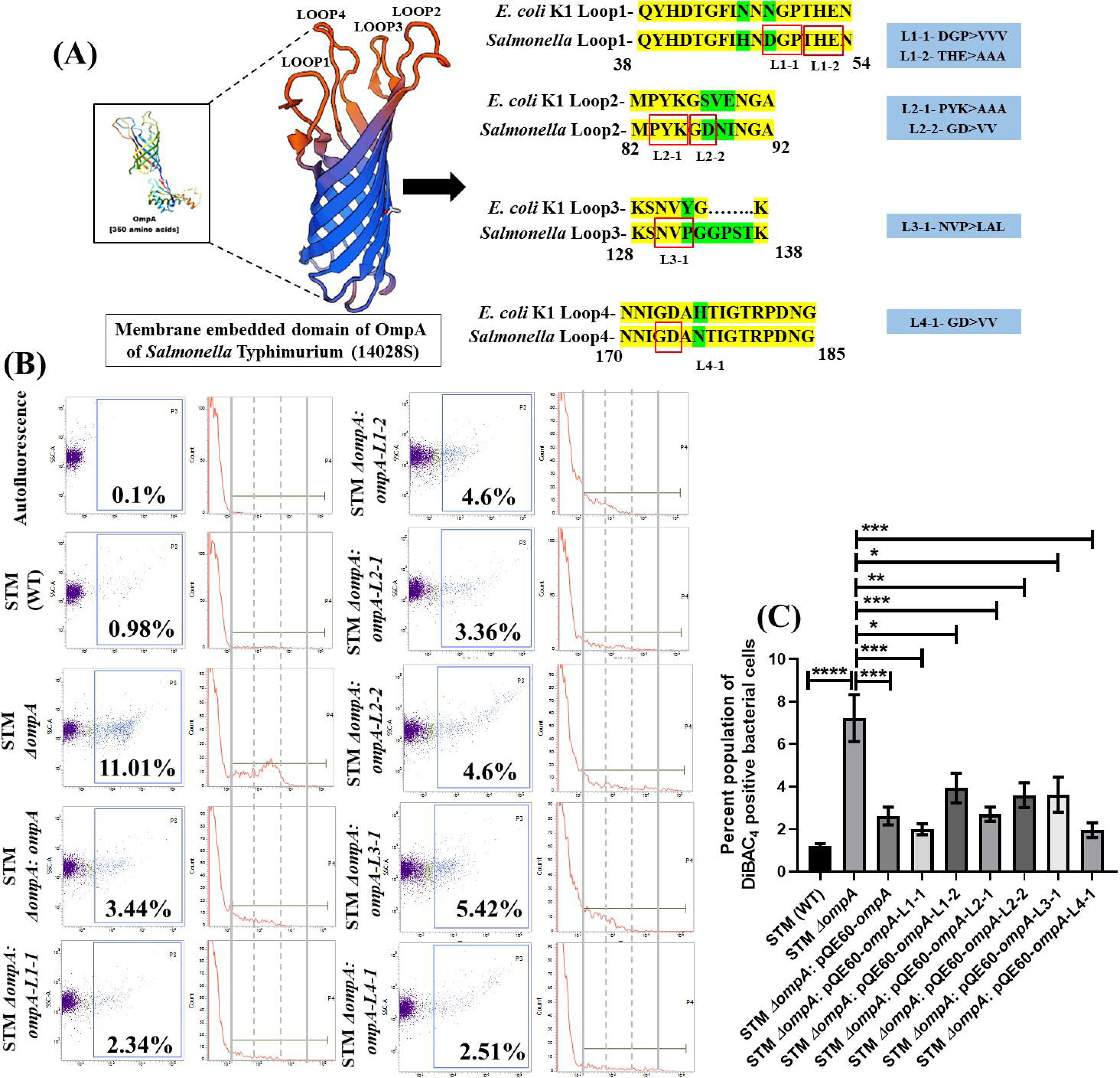
Introducing mutation in the extracellular loops of *Salmonella* Typhimurium OmpA by site-directed mutagenesis. (A) The outer membrane-embedded β barrel structure of *Salmonella* Typhimurium OmpA with extracellular loops (Loop1, Loop2, Loop3, and Loop4). Comparison between the extracellular loop sequences of *Escherichia coli* K1 and *Salmonella* Typhimurium. Two different mutations were introduced in loop1 (L1-L1-1 and L1-2) and loop2 (L2-L2-1 and L2-2) separately. Two distinct single mutations (L3-1 and L4-1) were introduced in loop3 (L3) and loop4 (L4). (B-C) Measurement of membrane porosity of STM (WT), *ΔompA*, *ΔompA*: pQE60-*ompA*, *ΔompA*: pQE60-*ompA-*L1-1, *ΔompA*: pQE60-*ompA-*L1-2, *ΔompA*: pQE60-*ompA-*L2-1, *ΔompA*: pQE60-*ompA-*L2-2, *ΔompA*: pQE60-*ompA-*L3-1, and *ΔompA*: pQE60-*ompA-*L4-1 in acidic F media (12 hours post-inoculation) using DiBAC4 (final concentration-1 µg/ mL) by flow cytometry. Unstained bacterial cells were used as control (Autofluorescence). Both dot plots (SSC-A vs. DiBAC4) and histograms (Count vs. DiBAC4) have been represented. The percent population of DiBAC4 positive bacterial cells has been represented here. Data are represented as mean ± SEM (n=5, N=2). ***(P) *< 0.05*, *(P) **< 0.005*, *(P) ***< 0.0005*, *(P) ****< 0.0001*, ns= non-significant**

### Inhibition of host autophagy using bafilomycin A restored the intracellular proliferation of *ompA* deficient strain of *Salmonella* Typhimurium

Earlier, it was found that when the macrophages were infected with STM *ΔompA,* there was an activation of the host autophagy machinery. This observation was strengthened by inhibiting the autophagy pathway using bafilomycin A1, a macrolide antibiotic isolated from *Streptomyces gresius*. Bafilomycin A1 inhibits the acidification of lysosomes by abrogating the vacuolar H^+^ ATPase pump [26]. In the absence of bafilomycin A1, we have seen that a significant population of STM *ΔompA* colocalizes with syntaxin17 (**Figure 2A and 2B**) and LC3B (**Figure 2C and 2D**) compared to the wild type bacteria, which is consistent with our previous findings. This suggests the formation of an active autophagosome which traps the mutant bacteria and restricts the infection within macrophages. When the cells were treated with bafilomycin A1 (50 nM), an impeded recruitment of syntaxin 17 and LC3B around STM *ΔompA* (**Figure 2A and 2C**) was found. Bafilomycin A1 treatment significantly reduced the percent colocalization between syntaxin 17 with STM *ΔompA* compared to the untreated macrophages infected with STM *ΔompA* (**Figure 2A and 2B**). On the other hand, a drastic abrogation in the formation of LC3B puncta was also noted around STM *ΔompA* under bafilomycin A1 treatment (**Figure 2C and 2D**). Normally, STM *ΔompA* has been found to be deficient in replication in murine macrophages compared to wild type bacteria (**Figure 2E**). When bafilomycin A1 was used, a significant recovery in the proliferation of STM *ΔompA* was observed. The acidification of SCV helps in the intracellular proliferation of wild type *Salmonella* by inducing SPI-2 gene expression. Bafilomycin A1 non-specifically inhibits the acidification of the cells’ acidic compartments, including both SCV and lysosomes. Hence it can slow down the growth of wild type bacteria residing within SCV, which can be considered as the reason behind the unaltered proliferation of STM (WT) upon bafilomycin A1 treatment (**Figure 2E**). Taken together, our data illustrate that *Salmonella* Typhimurium OmpA protects the bacteria against host autophagy machinery by improving the intracellular vacuolar life of the pathogen.

### STM *ΔompA* quits the SCV before the early stage of infection and remains in the cytosol of murine and human monocyte-derived macrophages

We further wanted to find out the time at which bacteria escape the SCV. To answer this question, both RAW264.7 and PMA activated U937 cells were infected with wild type, *ompA* mutant, and complemented strains of *Salmonella* and the vacuolar niche of the pathogen was investigated during early (2h post-infection) and middle (6h post-infection) stages of infection.

It was found that compared to the STM (WT), the colocalization of STM *ΔompA* with LAMP-1 is significantly less at 2 hours post-infection in RAW 264.7 cells (**Figure 3A and 3B**), stating that the knockout bacteria leave the vacuole even before the early stage of infection in macrophages. The reduced colocalization of the mutant bacteria with LAMP-1 was restored in the complemented strain (**Figure 3A and 3B**). Similarly, the poor colocalization of STM *ΔompA* with LAMP-1 at 6 hours post-infection suggests that majority of the bacteria that quit the vacuole before 2 hours remain in the cytosol during the middle stage of infection as well (**Figure 3C and 3D**). To inspect whether this phenotype is cell type-specific or not, the same experiment was carried out in PMA activated U937 cells. In accordance with our previous observation, it was found that STM *ΔompA* has a higher propensity towards abandoning the SCV before the early stage of infection than STM (WT) in U937 cells (**Figure S3A – S3F**). Hence it was decided to study the vacuolar niche of the pathogen during the immediate early phase of infection.

### STM *ΔompA* quits the SCV during the immediate early stage of infection in macrophages

The formation of SCV inside the host cell is a dynamic process. Immediately after entering the cells, the wild type *Salmonella* residing within early SCV attracts early endosome membrane markers such as early endosome antigen-1 (EEA-1), Rab5, and transferrin receptors, which are replaced within 20 to 40 minutes post-infection by late endosome membrane markers like- LAMP-1/2/3, Rab7, and V-ATPase [5, 27]. The lack of significant difference between the recruitment of EEA1 around STM (WT) and STM *ΔompA* at 15 minutes post-infection in activated U937 cells (**Figure 4A and 4B**) suggested that both the bacteria reside within the early SCV decorated with EEA1 during the immediate early stage of infection. In this experiment, PFA-fixed dead bacteria were used as a control. Taken together, this data also suggests that deletion of *ompA* from *Salmonella* does not hamper the SCV biogenesis during infection. Simultaneously, the colocalization of wild type and mutant bacteria with LAMP-1 was tested at 15-, 30- and 120 minutes post-infection in activated U937 (**Figure 4C-4F**) and RAW264.7 cells (**Figure 4G-4J**). At 15 minutes post-infection, STM *ΔompA* was found to be recruiting more LAMP-1 compared to the wild type bacteria in U937 (**Figure 4C and 4F**) and RAW264.7 (**Figure. 4G and 4J**) cells. With an increase in time (at 30- and 120 minutes post-infection), the wild type *Salmonella* was found to be acquiring more LAMP-1 in both the cells (**Figure 4D-4F and Figure 4H-4J**). In contrast, STM *ΔompA* was unable to retain the acquired LAMP-1 and started losing the SCV membrane. Taken together, our data demonstrated an essential role of outer membrane protein A of *Salmonella* to maintain a stable interaction with LAMP-1. Earlier, it was found that wild type *Salmonella* uses SPI-1 encoded virulent factor SipC (*Salmonella* invasion protein C) to acquire LAMP-1 from Golgi in a host syntaxin 6 dependent manners [28]. Our study revealed that STM *ΔompA* is unable to obtain LAMP-1 and escape the SCV. Hence, we hypothesized that STM *ΔompA* is deficient in producing SipC. To test this hypothesis, RAW264.7 cells were infected with wild type and *ompA* knockout strains of *Salmonella,* and the bacterial colocalization with SipC was studied. Surprisingly, no significant difference in the colocalization of SipC with intracellular STM (WT) and STM *ΔompA* (**Figure S4A and S4B**) was observed. Simultaneously, the expression of *sipC* transcripts from intracellular wild type and mutant bacteria was quantified (**Figure S4C**). There was no significant difference between the expression of *sipC* in wild type and *ompA* knockout strains of *Salmonella* proliferating in macrophages (**Figure S4C**), which is consistent with our previous findings. These data led us to conclude that OmpA plays a SipC independent role in maintaining a stable interaction of *Salmonella* with LAMP-1.

### *Salmonella* Typhimurium OmpA plays a critical role in maintaining host LAMP-1 (SCV) interaction during infection in macrophages

Earlier, we have shown an enhanced expression of *ompA* transcript in the wild type *Salmonella* Typhimurium growing intracellularly in murine macrophages at 9^th^ and 12^th^ hours post-infection, which suggested the requirement of OmpA for the intracellular survival of bacteria [15]. Our current study revealed that in the absence of OmpA, *Salmonella* could not stay inside the SCV. Hence, we hypothesized that outer membrane protein A plays a direct role in stabilizing the SCV membrane by retaining the LAMP-1 pool. To prove the interaction between *Salmonella* OmpA and host LAMP-1, the macrophages were infected with wild type bacteria, and the percent colocalization between OmpA and LAMP-1 during different time points was investigated. In RAW264.7 cells, the wild type bacteria use OmpA in maintaining a stable interaction with LAMP-1 at 2- and 6 hours post-infection (**Figure 5B and 5C**). This interaction was further increased during the late phase of infection (at 16 hours post-infection) (**Figure 5B and 5C**). In activated U937 cells, the interaction between OmpA with the LAMP-1 was found to be increasing significantly with time (**Figure 5D and 5E**). Taken together, our data suggested that indeed wild type *Salmonella* Typhimurium uses OmpA to retain the LAMP-1 around the SCV firmly. When OmpA is deleted, the bacteria could not hold the LAMP-1 pool and were gradually released into the cytosol of macrophages from the SCV. OmpA is embedded into the outer membrane of *Salmonella* Typhimurium with the help of its cylindrical structure consisting of eight anti-parallel β sheets. The β sheets are connected to each other by four periplasmic turns and four extracellular loops, which are exposed outside (**Figure 6A**), and likely to be interacting with LAMP-1. Hence, we decided to find out the role of these extracellular loops in establishing the interaction between the bacteria with LAMP-1. The *Salmonella* OmpA extracellular loop sequences were compared with *Escherichia coli* K1 (**Figure 6A**), another Gram-negative bacterial pathogen causing meningitis in neonates [29, 30]. Without hampering the membrane-embedded cylindrical structure of OmpA, the conserved and unique domains of the loops were altered by site-directed mutagenesis (**Figure 6A**), and six different variants, namely L1-1, L1-2, L2-1, L2-2, L3-1, L4-1, were generated. The mutated versions of the gene were expressed in the *ompA* knockout background of *Salmonella* and used for infection. The latest research from our group demonstrated that the depletion of OmpA increases the permeability of the bacterial outer membrane [15]. To validate the proper folding and localization of OmpA on the bacterial outer membrane despite receiving mutations in the extracellular loop regions, the outer membrane permeability of STM (WT), *ΔompA*, *ΔompA*: pQE60-*ompA*, *ΔompA*: pQE60-*ompA-*L1-1, *ΔompA*: pQE60-*ompA-*L1-2, *ΔompA*: pQE60-*ompA-*L2-1, *ΔompA*: pQE60-*ompA-*L2-2, *ΔompA*: pQE60-*ompA-*L3-1 and *ΔompA*: pQE60-*ompA-*L4-1 was checked by DiBAC_4_ staining (**Figure 6B and 6C**). In line with our expectations, an enhanced uptake of DiBAC_4_ was observed in STM *ΔompA* compared to the wild type and complemented strains of *Salmonella*. The complementation of the *ompA* knockout strain with mutated variants of *ompA* significantly reduced the entry of DiBAC_4_, suggesting a restoration of the outer membrane stability of the bacteria due to proper folding and localization of mutated OmpA (**Figure 6B and 6C**).

### Mutation in the extracellular loops of *Salmonella* Typhimurium OmpA hampers the stability of SCV but is not sufficient to target the bacteria to the lysosome

To find out the role of the extracellular loops of OmpA in maintaining the interaction of the bacteria with LAMP-1, the loop mutants of OmpA were used for infecting murine macrophages, and the recruitment of LAMP-1 was studied at 6- and 16 hours post-infection. Altering the aminoacid sequences in any one of the four extracellular loops of OmpA gave outcomes similar to STM *ΔompA*. Unlike the wild type and the complemented strains, all the loop mutants quit the vacuole at the 6^th^ hour post-infection (**Figure S5A**) and stayed in the cytosol of macrophages during the late phase of infection (16^th^ hour post-infection) as well (**Figure 7A**). The cytosolic localization of the loop mutants was confirmed by quantifying their percent colocalization with LAMP-1, which was comparable to the *ompA* knockout bacteria (**Figure S5B and 7B**). This suggests that the extracellular loops of *Salmonella* OmpA execute an important role in maintaining the stability of the SCV membrane by retaining LAMP-1.

**Figure 7.**
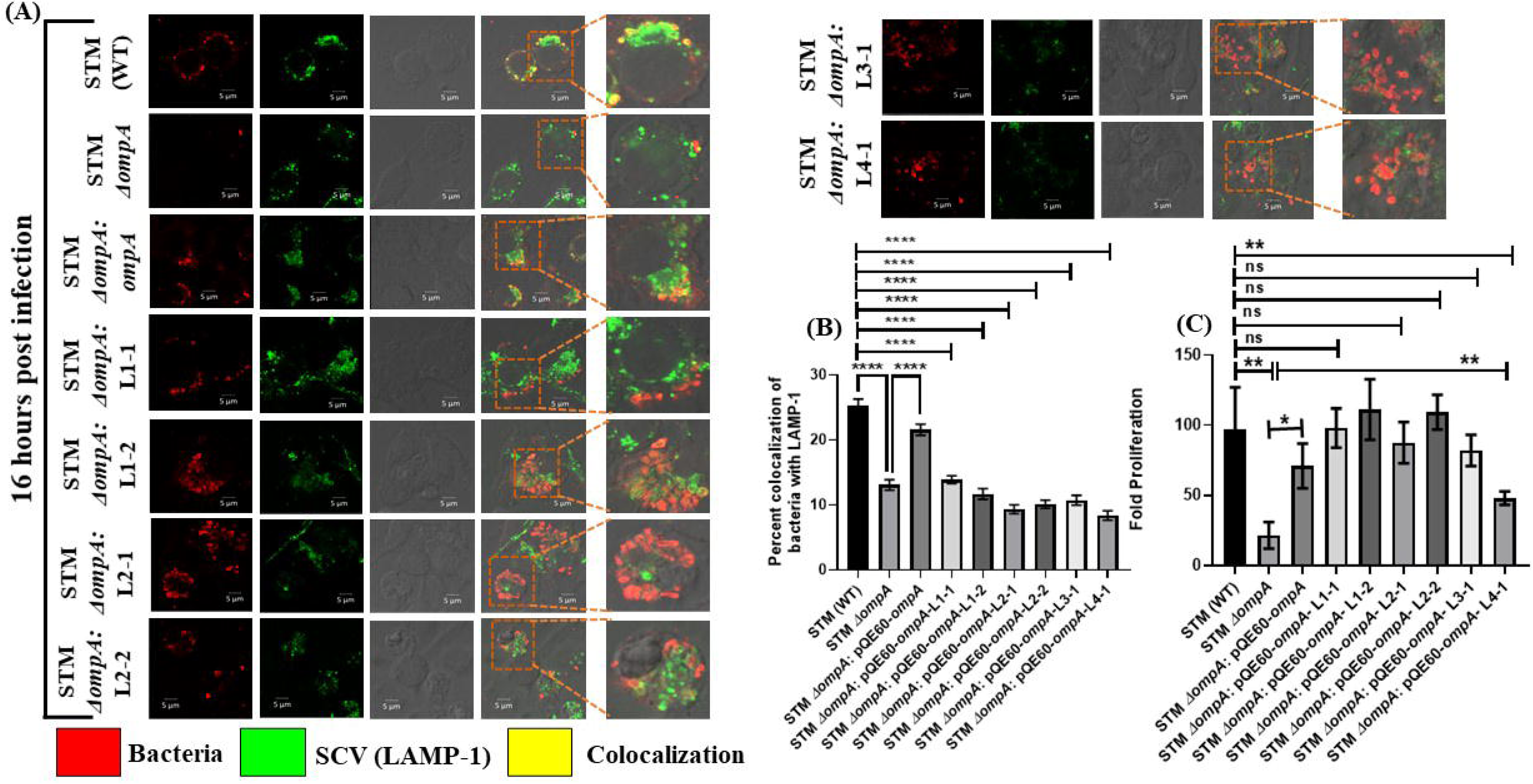
Mutation in the extracellular loops of *Salmonella* Typhimurium OmpA reduces the retention of LAMP-1 around the bacteria in murine macrophages. RAW264.7 cells were infected with STM (WT), *ΔompA*, *ΔompA*: pQE60-*ompA*, *ΔompA*: pQE60-*ompA-*L1-1, *ΔompA*: pQE60-*ompA-*L1-2, *ΔompA*: pQE60-*ompA-*L2-1, *ΔompA*: pQE60-*ompA-*L2-2, *ΔompA*: pQE60-*ompA-*L3-1 and *ΔompA*: pQE60-*ompA-*L4-1 at MOI of 20. Cells were fixed at (A) 16 hours (late phase) post-infection. Intracellular *Salmonella* was stained with rabbit-raised anti-*Salmonella* O primary antibody. LAMP-1 was labeled with rat-raised anti-mouse LAMP-1 primary antibody. The quantification of LAMP-1 recruitment on bacteria in RAW 264.7 cells at 16 hours post-infection has been represented in a graph. (B) During the late stage of infection (16 hours post-infection), the percent colocalization of bacteria with LAMP-1 was determined after analyzing 100 different microscopic fields from three independent experiments [n=100, N=3]. Scale bar = 5μm. (C) Intracellular survival of STM (WT), *ΔompA*, *ΔompA*: pQE60-*ompA*, *ΔompA*: pQE60-*ompA-*L1-1, *ΔompA*: pQE60-*ompA-*L1-2, *ΔompA*: pQE60-*ompA-*L2-1, *ΔompA*: pQE60-*ompA-*L2-2, *ΔompA*: pQE60-*ompA-* L3-1 and *ΔompA*: pQE60-*ompA-*L4-1 (MOI-10) in RAW264.7 cells (16 hours post-infection). The bacteria’s fold proliferation was calculated by normalizing the CFU at 16 hours to CFU at 2 hours (n=3, N=3). ***(P)* *< 0.05, *(P)* **< 0.005, *(P)* ***< 0.0005, *(P)* ****< 0.0001, ns= non-significant, (Student’s *t-*test).**

Bacteria were unable to hold the LAMP-1 around the SCV either in the absence of the entire OmpA (*ompA* knockout *Salmonella*) or due to the structural and functional ineffectiveness of the extracellular loops of OmpA. We further wanted to check the effect of the mutations in the loop region of OmpA on the intracellular survival of the bacteria. An intracellular survival assay was performed with the loop mutants in macrophages. Despite their cytosolic inhabitation, mutations in the loop region did not cast any impact on the intracellular survival of the bacteria. All the loop mutants were found to be surviving better than the STM *ΔompA* while infecting the macrophage (**Figure 7C**). Despite having mutations in the extracellular loop regions, the presence of intact OmpA on the outer membrane of the mutant bacteria might be the reason behind their better survival in the cytosol. We speculated that the cylindrical structure of OmpA, present in the outer membrane of the loop mutants, might be protecting the cytosolic bacteria from being targeted to the lysosome. To test our hypothesis, the colocalization of the loop mutants with the lysosomes was checked during the late phase of infection in murine macrophages (**Figure 8A and 8B**). Unlike the wild type and the complemented strains of *Salmonella*, STM *ΔompA* was found to be engulfed by the lysosomes during the late phase of infection in murine macrophages (**Figure 8A and 8B**). In accordance with our expectations, it was observed that the *ompA* variants having mutations in the extracellular loops are less prone to be captured by the lysosomes, which further explains their better survival within murine macrophages.

**Figure 8.**
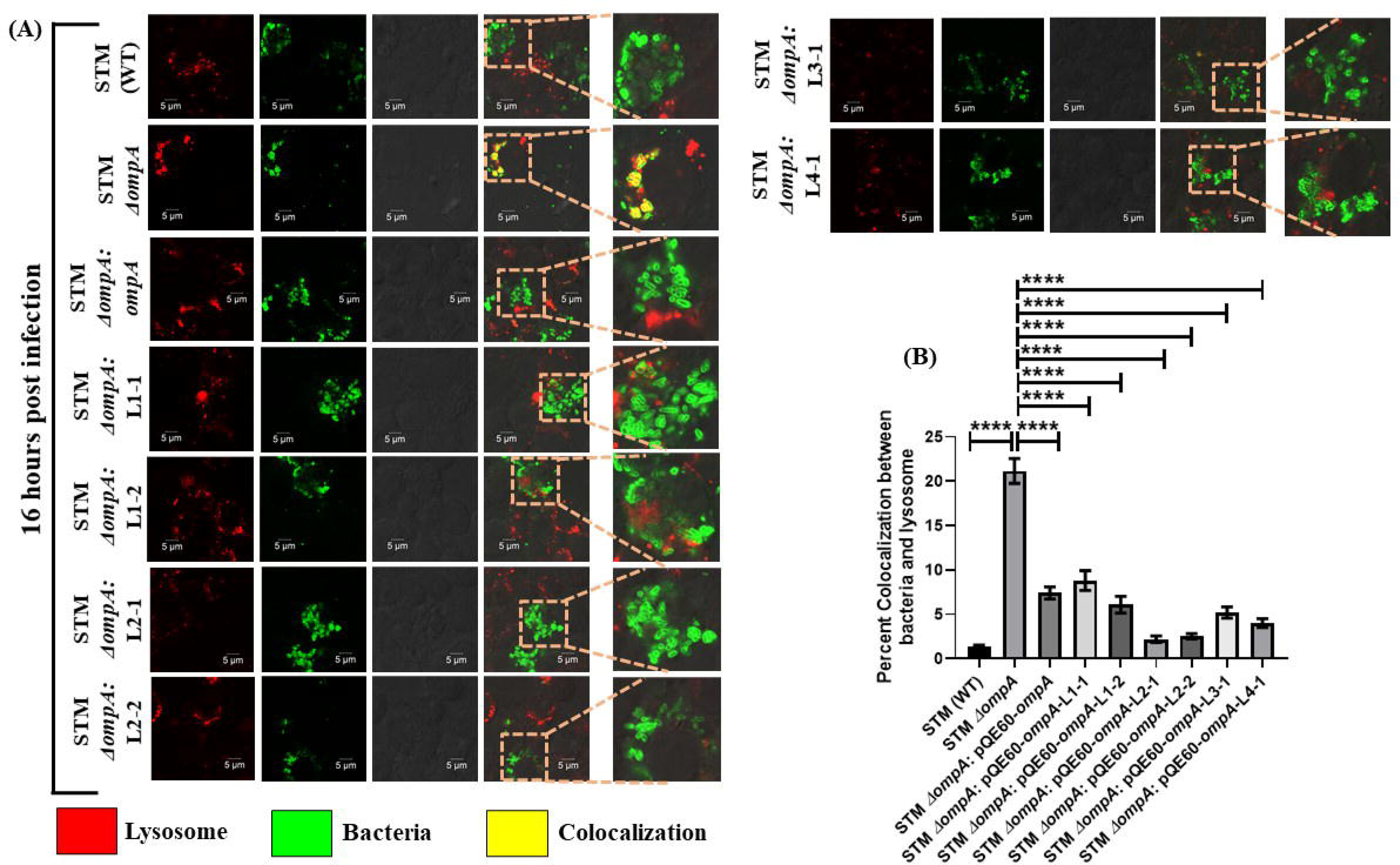
The mutation in the extracellular loops of OmpA is not sufficient to send the cytosolic population of *Salmonella* to the lysosomal degradation pathway. (A) RAW 264.7 cells were pre-treated with Texas red ovalbumin for 30 minutes and infected with STM (WT), *ΔompA*, *ΔompA*: pQE60-*ompA*, *ΔompA*: pQE60-*ompA-*L1-1, *ΔompA*: pQE60-*ompA-*L1-2, *ΔompA*: pQE60-*ompA-*L2-1, *ΔompA*: pQE60-*ompA-*L2-2, *ΔompA*: pQE60-*ompA-*L3-1 and *ΔompA*: pQE60-*ompA-*L4-1 respectively, at MOI of 20. Rabbit-raised anti-*Salmonella* O primary and anti-rabbit dylight 488 secondary antibodies were used to stain the intracellular bacteria. (B) The percent colocalization of bacteria with lysosome have been represented in the form of a graph. The percent colocalization between bacteria and texas red was determined after analyzing 100 microscopic stacks from two independent experiments (n=100, N=2). Scale bar = 5μm. ***(P)* ****< 0.0001, ns= non-significant, (Student’s *t-*test).**

**Figure 9.**
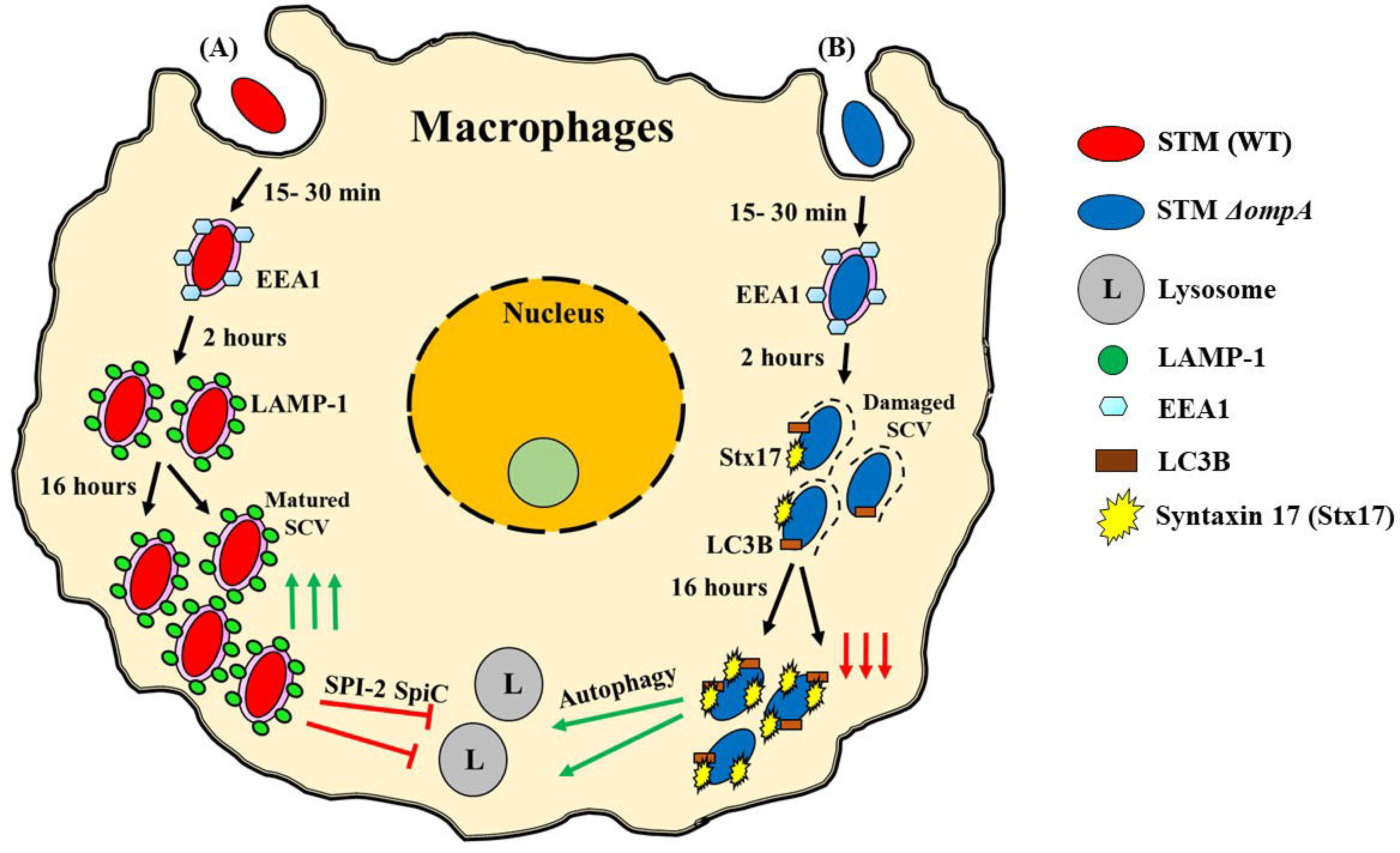
The hypothetical working model of activation of autophagy upon SCV damage by STM *ΔompA*. (A) After entering the host cell, STM (WT) stays inside the early endosomes. As time passes, EEA1 (early endosomal marker) is replaced with LAMP-1 (SCV marker). STM (WT) staying inside SCV can down-regulate lysosome biogenesis and suppress host autophagy machinery in SPI-2 encoded T3SS dependent manner. The down-regulation of lysosome biogenesis facilitates the successful proliferation of the bacteria inside macrophages. (B) Intracellular STM *ΔompA* is unable to retain LAMP-1 and comes into the cytosol after damaging SCV during the immediate early stage of infection. The extracellular loops of *Salmonella* OmpA play an essential role in maintaining the interaction between SCV and LAMP-1. The cytosolic population of STM *ΔompA* activates host autophagy machinery. After being colocalized with syntaxin 17 and LC3B, a significant fraction of cytosolic STM *ΔompA* is targeted to lysosomal degradation.

## Discussion

While infecting the host cell, *Salmonella* Typhimurium stays inside a modified compartment called *Salmonella* containing vacuole (SCV) [3, 4]. The low pH of the SCV triggers the activation of the PhoP/Q two-component system, which finally upregulates the expression of SPI-2 genes [31–33]. With the help of SPI-2 encoded T3SS2 and other virulent factors, *Salmonella* can inhibit the phagolysosome maturation and suppresses the lysosome biogenesis in the host cells [25, 34]. The SPI-2 genes of intracellular bacteria further impede the recruitment of iNOS and NADPH phagocytic oxidase around the SCV and ensure the successful proliferation of the bacteria within the host cells [35, 36]. The formation of SCV is a dynamic and complex process that employs a wide array of host and bacterial proteins. Immediately after entering the host cells, the bacteria-containing vacuole recruits the early endosome membrane markers such as EEA1, transferrin receptors, and Rab5. With time, the early endosomal proteins are replaced with late endosome membrane markers such as LAMP-1, LAMP-2, Rab7, etc. [4]. *Salmonella* uses SPI-1 and SPI-2 encoded virulent factors to regulate the biogenesis and stability of the SCV. SPI-1 encoded virulent factor SopB phosphatase can reduce the membrane charge of SCV and prevent the fusion of lysosomes with SCV [11]. *Salmonella* employs SPI-2 encoded protein SpiC to inhibit the fusion of lysosomes with SCV [34]. Another SPI-2 protein, SifA, forms *Salmonella*-induced filaments in epithelial cells to maintain the integrity of SCV by downregulating the recruitment of kinesins [37, 38]. SifA interacts with Rab9 to block the retrograde transport of mannose-6-phosphate receptors and MPR hydrolases to the Golgi apparatus, which ultimately inhibits lysosomal fusion with SCV [39]. Tampering (by point mutation or deletion) the host and bacterial effectors that control the stability of SCV create a tremendous impact on the intracellular fate of the pathogen. Introducing point mutation in host Rab7 or deleting *sifA* from *Salmonella* released the bacteria into the host cytosol from SCV, which makes the bacteria hyper-virulent in epithelial cells and replication-deficient in macrophages [7, 8]. On the contrary, deletion of SPI-2 gene *sseJ* in *sifA* null background further restored the vacuolar status of the pathogen [9]. However, very little is known about the contribution of non-SPI virulent genes of *Salmonella* on the biogenesis and integrity of SCV.

Our study revealed a novel role of *Salmonella* Typhimurium outer membrane protein A (OmpA) in maintaining the stability of SCV. Outer membrane protein A is an outer membrane-bound porin of *Salmonella* that protects the pathogen from oxidative and nitrosative stress [13, 15]. The deletion of OmpA compromises the intracellular vacuolar status of the bacteria. STM *ΔompA* was unable to recruit LAMP-1 around the SCV and gradually was released into the host cytosol. The SCV quitting phenomenon of the pathogen was reversed when the gene was complemented in the knockout bacteria. Earlier, our group has shown that this cytosolic population of the bacteria becomes hyper-proliferative inside the epithelial cells, and their intracellular proliferation was significantly attenuated in macrophages [15]. Our study further depicted that STM *ΔompA* is unable to recruit T3SS2 translocon proteins (SseC and SseD) on its surface and is deficient in producing SPI-2 effectors such as SsaV and SpiC [15]. Available literature suggests that *Salmonella* deficient in making active T3SS2 and with damaged SCV cannot suppress host autophagy machinery. Hence, we wanted to investigate the autophagy-inducing ability of the cytosolic population of STM *ΔompA*. Compared to the wild type bacteria, which stay within LAMP-1 decorated SCV, the cytosolic population of STM *ΔompA* colocalizes more with syntaxin 17 and LC3B. The inhibition of autophagy using bafilomycin A1 not only reduced the recruitment of autophagy markers (syntaxin 17 and LC3B) on STM *ΔompA* but also improved their intracellular life. It was speculated that the activation of the autophagy pathway would recruit lysosomes around the mutant bacteria. By Texas red ovalbumin pulse-chase experiment, an enhanced colocalization of lysosomes with STM *ΔompA* was observed in macrophages. When STM (WT): *LLO* (which stays in the macrophage cytosol with intact OmpA on its outer membrane) was used for infection, a reduced colocalization between the lysosome and bacteria was observed, suggesting that OmpA protects the cytosolic bacteria from being targeted to lysosomes. This proves that the syntaxin 17 and LC3B recruited around the cytosolic bacteria lacking OmpA can target them into the lysosomes. We confirmed this result by measuring the enhanced activity of lysosomal acid phosphatases in the cells infected with STM *ΔompA*. Earlier, it has been reported that *Salmonella* uses SPI-2 encoded virulent factor SpiC to inhibit phagosome-lysosome fusion [34]. STM *ΔompA* has been found to be deficient in producing SpiC, which might be the reason behind the enhanced colocalization between the bacteria with host lysosomes.

We further wanted to check at what time the bacteria quit the vacuole in the absence of OmpA. A significant population of STM *ΔompA* escapes the SCV during the early stage of infection in macrophages. Once this bacterial population left the SCV, it remained in the cytosol during the rest of the infection. This result helped us speculate that the departure of STM *ΔompA* from SCV is happening even before the early stage of infection. Hence the recruitment of LAMP-1 around the bacteria was examined at 15-, 30- and 120 minutes post-infection. It was found that, immediately after entering the macrophages, both the wild type and the mutant bacteria stay within EEA1^+^ early SCV, suggesting uninterrupted biogenesis of SCV. Moreover, at this stage of infection (15 minutes p.i.), STM *ΔompA* recruits more LAMP-1 than wild type bacteria. With an increase in time, STM *ΔompA* was unable to retain the LAMP-1 pool and gradually lost the SCV membrane, unlike the wild type bacteria. Taken together, it was concluded that without changing the biogenesis of SCV, the absence of OmpA in *Salmonella* only hampers the integrity of SCV by restricting the recruitment of LAMP-1. Wild type *Salmonella* uses SPI-1 effector protein SipC to recruit LAMP-1 from Golgi in host syntaxin 6 dependent manners [28]. As it was observed that STM *ΔompA* is unable to hold the LAMP-1 pool, we speculated that STM *ΔompA* is deficient in producing SipC. The comparable colocalization of SipC between the intracellular wild type and mutant bacteria proved that the OmpA deletion mutant works in a SipC independent manner. This conclusion was supported by measuring the unaltered *sipC* transcript level from intracellular wild type and mutant bacteria. This result motivated us to hypothesize that OmpA plays an important structural role in maintaining the interaction of LAMP-1 with the bacteria confined inside the SCV. The direct interaction between the LAMP-1 and the OmpA of wild type bacteria was measured to verify the hypothesis. The intracellular wild type *Salmonella* was stained with anti-*Salmonella* OmpA antibody, and the recruitment of LAMP-1 around the bacteria was estimated. We have seen that *Salmonella* Typhimurium OmpA maintains a stable interaction with the host LAMP-1 during the early and middle stages of infection in macrophages, which further increases during the late phase of infection.

Outer membrane protein A is embedded into the bacterial outer membrane with the help of its cylindrical β barrel structure. The anti-parallel β sheets that constitute the wall of this barrel structure are connected by four extracellular loops (L1, L2, L3, and L4), which are exposed outside. The externally exposed extracellular loops were thought to maintain the integrity of SCV by directly interacting with host LAMP-1. Our speculation was validated by introducing mutations in the loop regions of *Salmonella* OmpA by site-directed mutagenesis. We have performed a comparative study on the amino acid sequences of the extracellular loops of OmpA between *Salmonella* Typhimurium and *Escherichia coli* K1 [40]. The role of the extracellular loops of *E. coli* K1 OmpA has already been deciphered. The introduction of mutations in the loop regions of *E. coli* K1 OmpA made the bacteria proliferation deficient in immune cells and reduced their ability to cause meningitis in a neonatal mouse model [29, 41]. The unique and the conserved sequences of the extracellular loops of *Salmonella* OmpA were targeted to make them deformed structurally and functionally. The amino acid sequences of the loop region were changed to nonpolar amino acids like-alanine, leucine, and valine. It was found that each of these individual loop mutants plays a critical role in maintaining the interaction with LAMP-1 during the middle and late stages of infection in murine macrophages. All the OmpA loop mutants came into the cytosol of macrophages after abandoning the SCV. Surprisingly, it was observed that the mutations in the loop regions do not hamper the intracellular proliferation of the bacteria within the cytosol of macrophages. To find out the reason behind the better survival of the *Salmonella* loop mutants in murine macrophages, their colocalization with the lysosomal compartments was examined. The reduced percent colocalization of the loop mutants with Texas red ovalbumin suggested poor recruitment of lysosomes on the cytosolic niche of *Salmonella* Typhimurium *ompA* variants. In the previous study, we have proved that the presence of intact OmpA maintains the integrity of the bacterial outer membrane and protects it from nitrosative stress. The final observation from this study led us to conclude that despite mutations in the extracellular loops, the intact OmpA present on the bacterial outer membrane could protect the cytosolic bacteria from lysosomal degradation in murine macrophages.

Altogether, our study provides an OmpA dependent novel mechanism used by *Salmonella* Typhimurium to maintain the stability and integrity of the SCV. *Salmonella* uses the extracellular loops of OmpA to retain the LAMP-1 pool around the SCV. In the absence of OmpA, the bacteria fail to hold the LAMP-1 and come into the cytosol after quitting the SCV. The cytosolic bacteria lacking OmpA further activate the host autophagy machinery. They are unable to prevent the maturation of the phagolysosome, which further leads to the clearance of the bacteria by the lysosomal degradation pathway.

## Materials and methods

### Bacterial strains, media, and culture conditions

The wild type (WT) bacteria *Salmonella enterica* serovar Typhimurium strain 14028S used in this study was a generous gift from Professor Michael Hensel, Max Von Pettenkofer-Institute for Hygiene and Medizinische Mikrobiologie, Germany. The bacterial strains were revived from glycerol stock (stored in -80^0^C) and plated either only on LB agar (purchased from HiMedia) (for the wild type *Salmonella*) or LB agar along with appropriate antibiotics like-kanamycin (50 μg/mL) (for the *ompA* knockout strains), ampicillin (50 μg/mL) (for the wild type *Salmonella* expressing mCherry/ RFP, GFP, and LLO), and kanamycin and ampicillin together (both 50 μg/mL), (for the complemented, loop mutants, mCherry and GFP expressing *ompA* knockout strains). *Salmonella*-*Shigella* agar was used for plate cell lysates/ cell suspensions to calculate the bacterial burden in infected cell lines. The complete list of strains and plasmids has been listed below. (Description in Table-4.1) Dead bacteria used in several experiments were produced from viable wild type bacteria by treating the bacteria with 3.5% paraformaldehyde for 30 minutes.

**Table 4.1.**
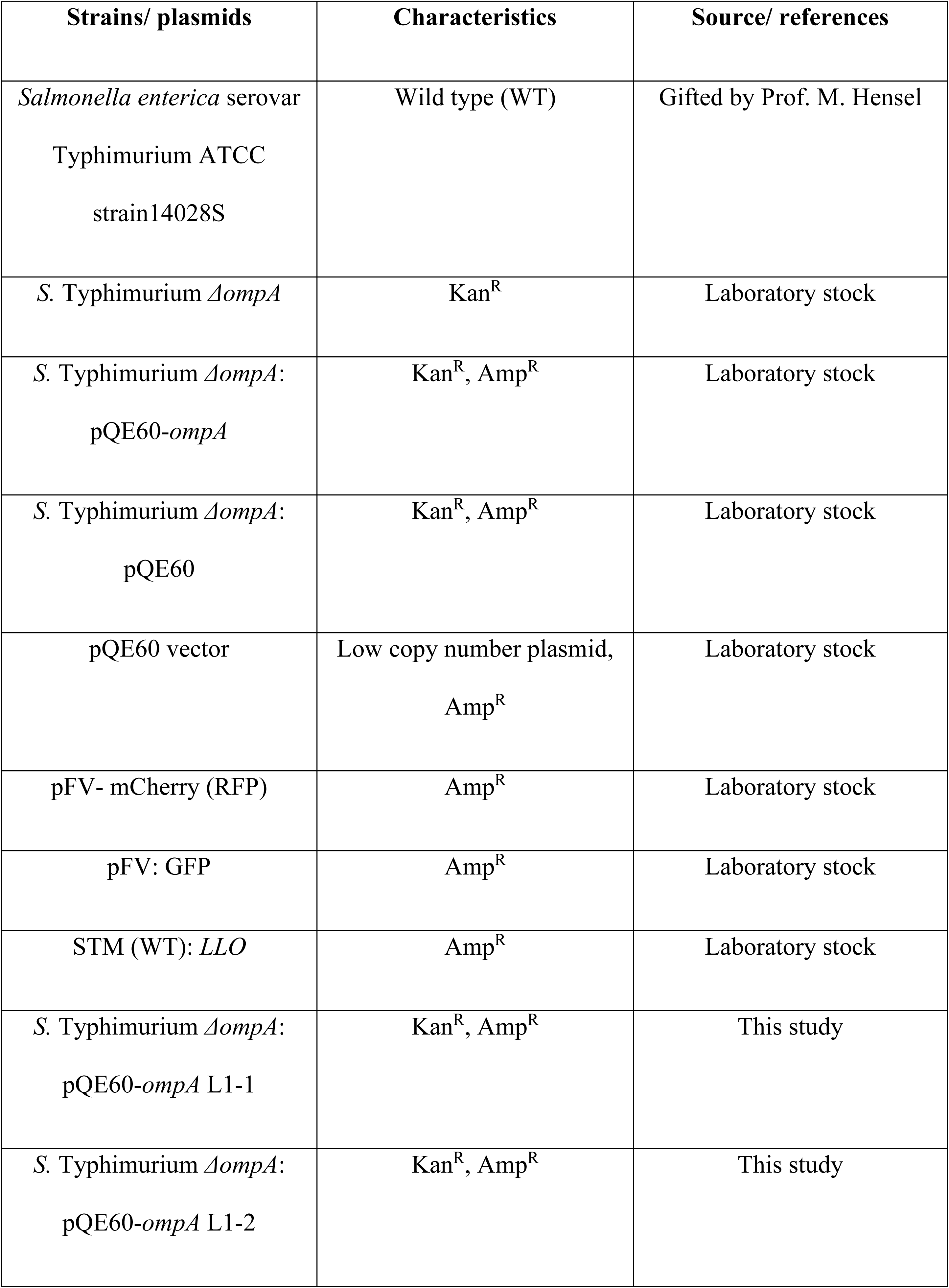
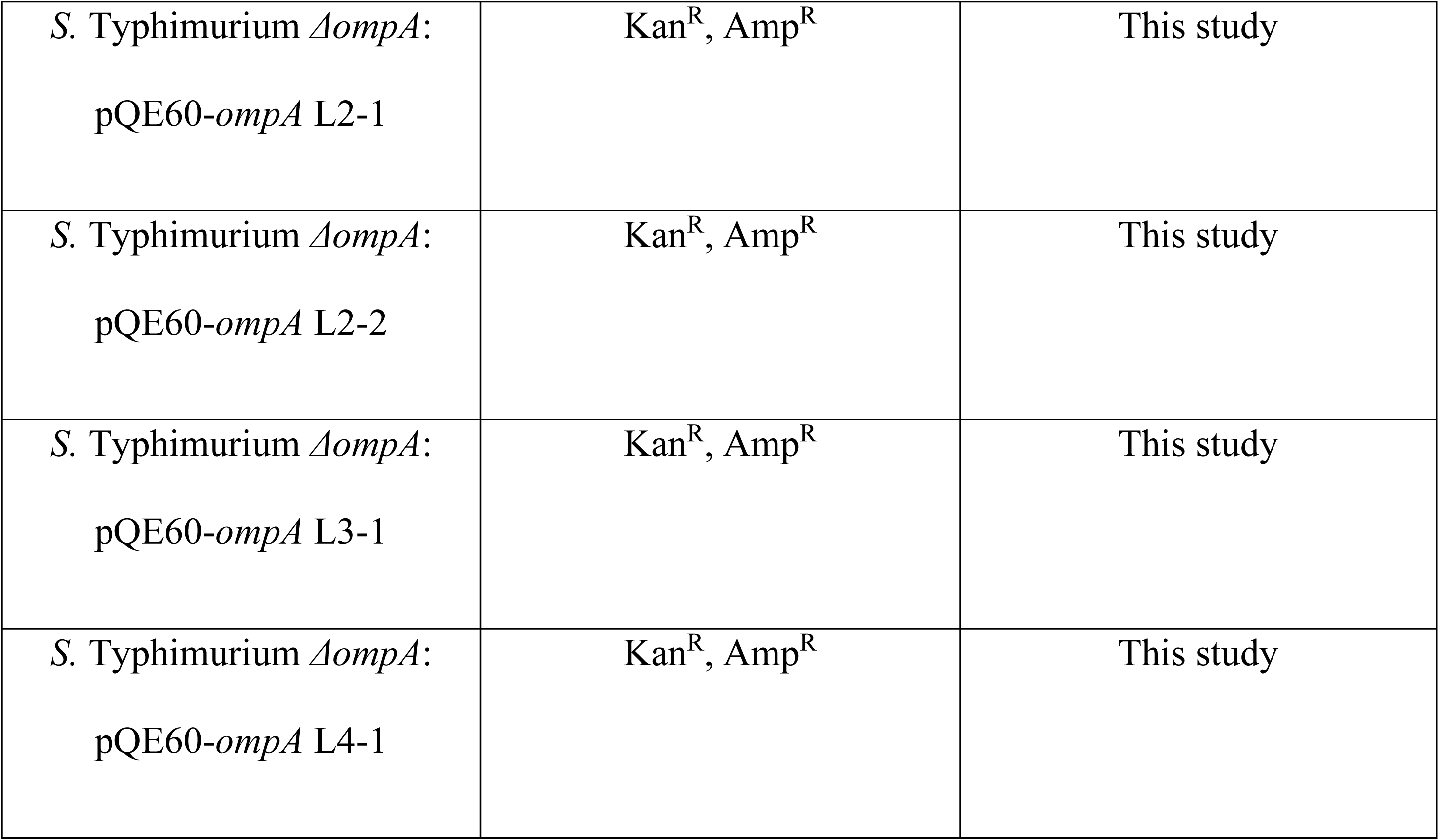
**Strains and plasmids used in this study**

| Strains/ plasmids | Characteristics | Source/ references |
| --- | --- | --- |
| *Salmonella enterica* serovar<br>Typhimurium ATCC<br>strain14028S | Wild type (WT) | Gifted by Prof. M. Hensel |
| *S. Typhimurium* $\Delta ompA$ | Kan<sup>R</sup> | Laboratory stock |
| *S. Typhimurium* $\Delta ompA$:<br>pQE60-*ompA* | Kan<sup>R</sup>, Amp<sup>R</sup> | Laboratory stock |
| *S. Typhimurium* $\Delta ompA$:<br>pQE60 | Kan<sup>R</sup>, Amp<sup>R</sup> | Laboratory stock |
| pQE60 vector | Low copy number plasmid,<br>Amp<sup>R</sup> | Laboratory stock |
| pFV- mCherry (RFP) | Amp<sup>R</sup> | Laboratory stock |
| pFV: GFP | Amp<sup>R</sup> | Laboratory stock |
| STM (WT): *LLO* | Amp<sup>R</sup> | Laboratory stock |
| *S. Typhimurium* $\Delta ompA$:<br>pQE60-*ompA* L1-1 | Kan<sup>R</sup>, Amp<sup>R</sup> | This study |
| *S. Typhimurium* $\Delta ompA$:<br>pQE60-*ompA* L1-2 | Kan<sup>R</sup>, Amp<sup>R</sup> | This study |
| *S. Typhimurium* $\Delta ompA$:<br>pQE60-*ompA* L2-1 | Kan<sup>R</sup>, Amp<sup>R</sup> | This study |
| *S. Typhimurium* $\Delta ompA$:<br>pQE60-*ompA* L2-2 | Kan<sup>R</sup>, Amp<sup>R</sup> | This study |
| *S. Typhimurium* $\Delta ompA$:<br>pQE60-*ompA* L3-1 | Kan<sup>R</sup>, Amp<sup>R</sup> | This study |
| *S. Typhimurium* $\Delta ompA$:<br>pQE60-*ompA* L4-1 | Kan<sup>R</sup>, Amp<sup>R</sup> | This study |

### Eukaryotic cell lines and growth conditions

The murine macrophage-like cell lines RAW 264.7 used in this study were maintained in Dulbecco’s Modified Eagle’s Media (Sigma-Aldrich) supplemented with 10% FCS (Fetal calf serum, Gibco) at 37^0^C temperature in the presence of 5% CO_2_. Human monocyte cell line U937 cells were maintained in Roswell Park Memorial Institute 1640 media (Sigma-Aldrich) supplemented with 10% FCS (Fetal calf serum, Gibco). Phorbol Myristate Acetate (Sigma-Aldrich) (concentration- 20 ng/ mL) was used for the activation of U937 cells for 24 hours at 37^0^C temperature in the presence of 5% CO_2_, followed by the replacement of the media carrying PMA with normal RPMI supplemented with 10% FCS and further incubating the cells for 24 hours before starting the experiments.

### RNA isolation from intracellular bacteria and RT PCR

The RAW264.7 cells were infected with STM (WT) and *ΔompA* at MOI of 50. 12hours post-infection, the infected macrophages were lysed with TRIzol reagent (RNAiso Plus, Takara) and stored at -80^0^C overnight. The lysed supernatants were further subjected to chloroform extraction followed by precipitation of total RNA by adding an equal volume of isopropanol. The pellet was washed with 70% RNA-grade ethanol, air-dried, and suspended in 20 μL of DEPC water. The RNA concentration was estimated in nano-drop and run on 1.5% agarose gel to assess RNA quality. To make cDNA, 3 μg of RNA sample was subjected to DNase treatment (Thermo Fischer Scientific) at 37^0^C for two hours. The reaction was stopped by adding 5mM Na_2_EDTA (Thermo Fischer Scientific), followed by heating the sample at 65^0^C for 10 min. The samples were incubated with random hexamers at 65^0^C for 10 minutes and then supplemented with 5X RT buffer, RT enzyme, dNTPs, and DEPC treated water at 42^0^C for an hour. Quantitative real-time PCR was done using SYBR/ TB Green RT PCR kit (Takara Bio) in Bio-Rad real-time PCR detection system, and the expression level of target genes was measured using *Salmonella* Typhimurium *sipC* specific RT primers (Table-4.2). 16S rRNA transcript level was used to normalize the expression levels of the target genes.

**Table 4.2.**
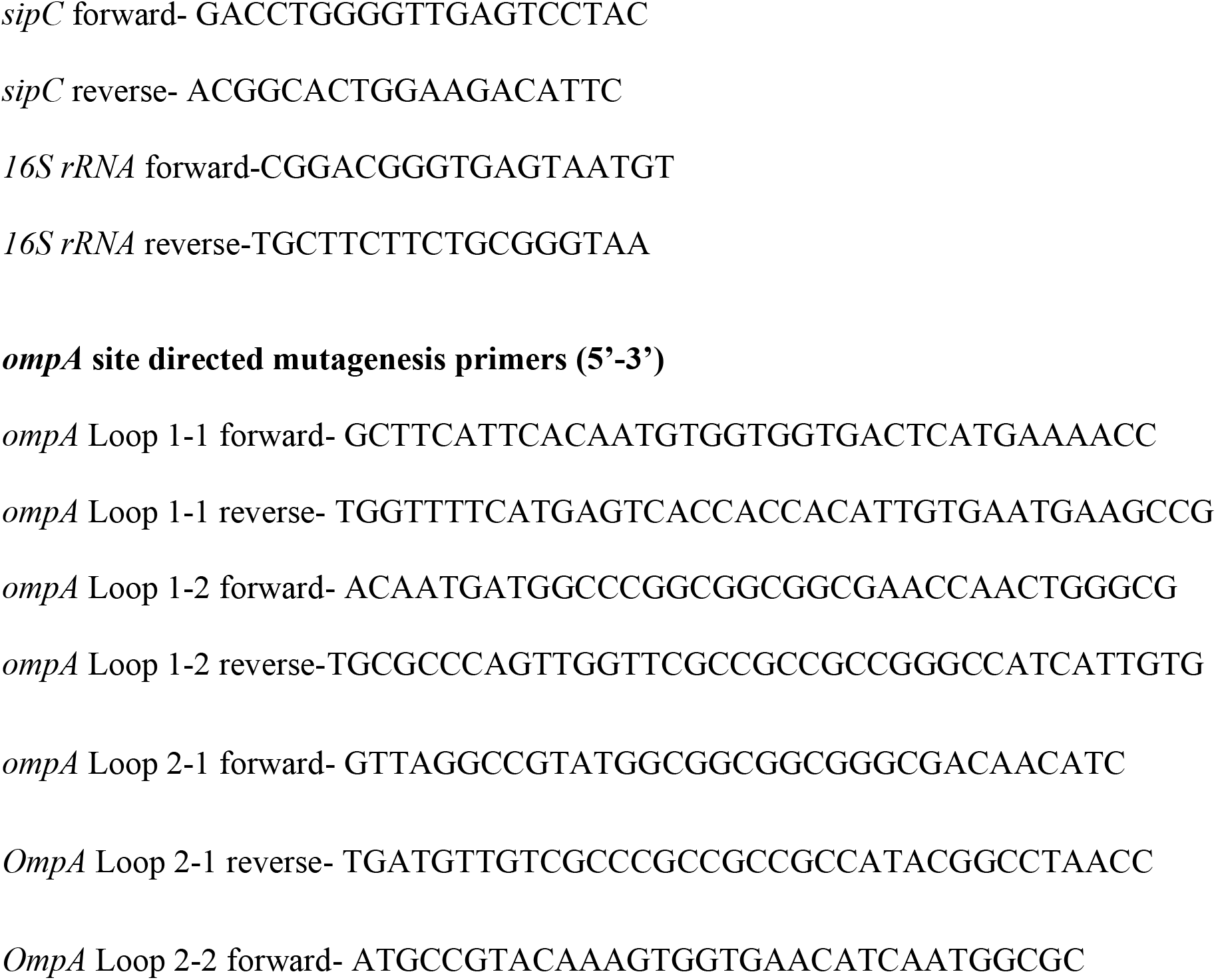
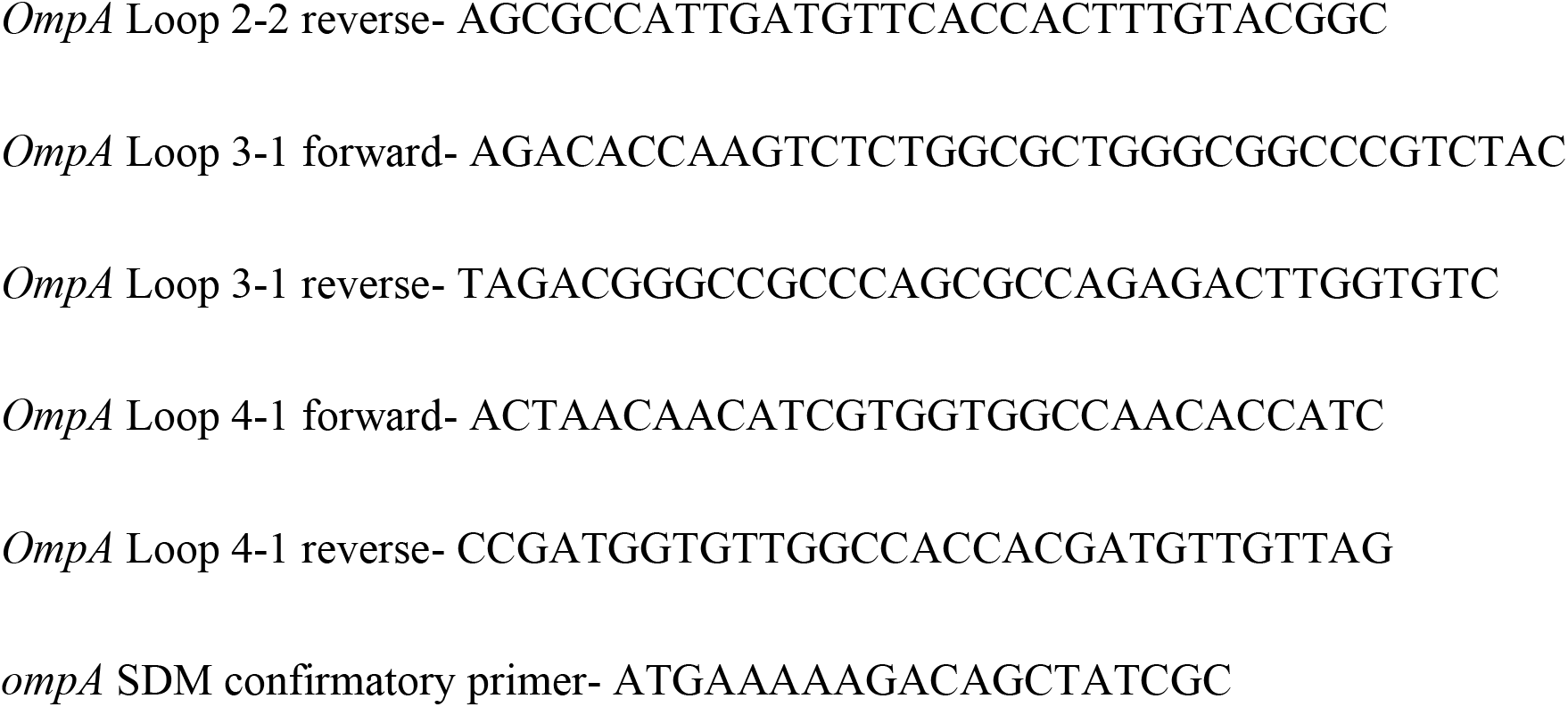
**Primers used in this study (5’-3’)**

### Intracellular proliferation assay

RAW264.7 (1.5 to 2 X 10^5^ cells seeded per well) were infected with STM (WT), *ΔompA, ΔompA*: pQE60-*ompA*, *ΔompA*:pQE60-*ompA* L1-1, *ΔompA*:pQE60-*ompA* L1-2, *ΔompA*:pQE60-*ompA* L2-1, *ΔompA*:pQE60-*ompA* L2-2, *ΔompA*:pQE60-*ompA* L3-1, and *ΔompA*:pQE60-*ompA* L4-1 at MOI of 10. After centrifuging the cells at 800 rpm for 5 minutes, the infected cells were incubated at 37^0^C temperature in the presence of 5% CO_2_ for 25 minutes. Next, the cells were washed thrice with PBS to remove all the unattached extracellular bacteria and subjected to 100 μg/ mL concentration of gentamicin treatment for 1 hour. This was followed by washing the cells with sterile PBS and subjecting them to a 25 μg/ mL concentration of gentamicin treatment till the lysis. The cells were lysed with 0.1% triton-X- 100 at 2 hours and 16 hours post-infection. The lysates were plated on *Salmonella- Shigella* Agar, and the corresponding CFU at 2 hours and 16 hours were determined. The intracellular proliferation of bacteria (Fold proliferation) was determined using a simple formula-

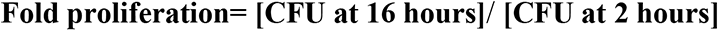

In some sets of experiments, the fold proliferation of STM (WT) and *ΔompA* in the macrophages (RAW 264.7) was measured in the presence of autophagy inhibitor bafilomycin A (50 nM). Bafilomycin A was added to the cells infected with STM (WT) and *ΔompA* along with 25 μg/ mL of gentamycin solution and incubated till the lysis. As usual, the cells were lysed with 0.1% triton-X-100 at 2 hours and 16 hours post-infection. The lysates were plated on *Salmonella- Shigella* Agar, and the corresponding CFU at 2 hours and 16 hours were calculated to determine Fold proliferation.

### Confocal microscopy

RAW 264.7 or U937 cells were seeded at a 1.5 to 2 X 10^5^ cells density per sterile glass coverslips. U937 cells were activated using PMA (as mentioned earlier in 4.2.2). The cells were infected with appropriate bacterial strains at MOI 20. The cells were washed thrice with PBS and fixed with 3.5% paraformaldehyde for 15 minutes at indicated time points post-infection. The cells were first incubated with specific primary antibody raised against- wild type *Salmonella* Typhimurium (rabbit raised anti- *Salmonella* O antigen), mouse lysosome-associated membrane protein-1 (LAMP-1) (rat raised anti-mouse LAMP-1), human LAMP-1 (mouse raised anti-human LAMP-1), mouse LC3B (rabbit raised anti-mouse LC3B), mouse syntaxin 17 (rabbit raised anti-mouse syntaxin 17), *Salmonella* Typhimurium SipC (mouse raised anti-*Salmonella* SipC), *S.* Typhimurium OmpA (rabbit raised anti-*Salmonella* OmpA antibody) and human EEA1 (mouse raised anti-human EEA1) as per the requirements of experiments. The primary antibodies were diluted in 2.5% BSA and 0.01% saponin (dilution 1: 100, duration 6 to 8 hours at 4^0^C temperature). This was followed by incubating the cells with appropriate secondary antibodies conjugated with fluorophores (dylight 488, alexa fluor 647, and cy3) (dilution 1: 200, duration 1 hours at room temperature). The coverslips were mounted with anti-fade reagent and fixed on a glass slide with transparent nail paint. Samples were imaged by confocal laser scanning microscopy (Zeiss LSM 710) using a 63X oil immersion objective lens. The images were analyzed with ZEN Black 2009 software provided by Zeiss.

### Texas red ovalbumin pulse-chase experiment

1.5 to 2 X 10^5^ RAW 264.7 cells were seeded on the top of sterile glass coverslips in the wells of a 24 well plate. The cells were fed with a 50 µg/ mL concentration of Texas red ovalbumin (resuspended in DMEM media) for 30 minutes at 37^0^C in the presence of 5% CO_2_. After this, the labeling media was removed, and the cells were washed with sterile PBS. The cells were further incubated for 30 minutes with fresh DMEM media and infected with overnight grown 10- 12 hours old stationary phase culture of STM (WT), STM (WT): *GFP*, *ΔompA*: *GFP,* and STM (WT): *LLO* at MOI of 20. PFA-fixed dead bacteria were used for infection as control at MOI of 25. The cells were washed thrice with PBS and fixed with 3.5% paraformaldehyde for 15 minutes at indicated time points post-infection. Rabbit-raised anti-*Salmonella* primary antibody stained the STM (WT): *LLO* and PFA fixed dead bacteria. The coverslips were mounted with anti-fade reagent and fixed on a glass slide with transparent nail paint to visualize the lysosomal arrangements in the infected macrophage cells. Samples were imaged by confocal laser scanning microscopy (Zeiss LSM 710) using a 63X oil immersion objective lens. The images were analyzed with ZEN Black 2009 software provided by Zeiss.

### Acid phosphatase assay

The protocol of acid phosphatase assay has been followed as mentioned earlier [25, 42, 43]. 1.5 to 2 X 10^5^ RAW 264.7 cells were seeded into the wells of a 24 well plate and infected with overnight grown stationary phase culture of STM (WT), *ΔompA, ΔompA*: pQE60-*ompA*, *ΔompA*: pQE60, and STM (WT): *LLO* at MOI of 10. PFA-fixed dead bacteria were used as a control for the infection (MOI-20). The infected cells were incubated for 12 hours under gentamycin treatment (as mentioned earlier). At the end of the incubation period, the cells were washed with PBS and incubated for 4 hours at 37^0^C with a buffer containing a 0.1 (M) sodium acetate of pH= 5, 0.1% triton-X-100, 5mM of *p-*nitrophenyl phosphate (pNPP). The absorbance of the supernatant was measured at 405 nm using a microplate reader. The non-enzymatic hydrolysis of pNPP (negligible) was measured media control without macrophage cells.

### Generation of extracellular loop mutants of *ompA* by site-directed mutagenesis

The protocol for site-directed mutagenesis was followed, as mentioned earlier [44]. Primer pairs (from Table-4.2) carrying mutations in the desired locations of the four extracellular loops of *Salmonella* Typhimurium OmpA (Figure 4.12.B) were used to amplify the pQE60- *ompA* recombinant plasmid (Size- 4.5 kb) using Phusion high fidelity DNA polymerase (NEB). The reaction mixture was heated at 95^0^C for 10 minutes for the plasmid’s denaturation, followed by 35 amplification cycles at 95^0^C for 30 seconds, 54^0^C for 60 seconds, and 72^0^C for 2 minutes 30 seconds with a final extension time of 10 minutes. The amplified PCR product was then digested with DpnI and transformed into *E. coli* TG1. The transformants were then selected on ampicillin plates. The plasmids were isolated from the transformant colonies, and the mutations were verified by sequencing. Once the confirmation was done, the recombinant plasmids carrying desired mutations were transformed into STM *ΔompA* by electroporation to create specific loop mutant *Salmonella*. These loop mutants were used for infecting RAW264.7 cells.

### Measurement of outer membrane porosity of *Salmonella* Typhimurium OmpA loop mutants

Outer membrane porosity of STM (WT), *ΔompA*, *ΔompA*: pQE60-*ompA*, *ΔompA*: pQE60- *ompA-*L1-1, *ΔompA*: pQE60-*ompA-*L1-2, *ΔompA*: pQE60-*ompA-*L2-1, *ΔompA*: pQE60-*ompA-* L2-2, *ΔompA*: pQE60-*ompA-*L3-1 and *ΔompA*: pQE60-*ompA-*L4-1 grown in low magnesium acidic F medium (pH= 5.4) for 12 hours was measured using a dye called bis-(1,3-dibutyl barbituric acid)-trimethylene oxonol (Invitrogen) (DiBAC4). Briefly, 4.5 X 10^7^ CFU of each bacterial strain was incubated with 1 µg/ml of DiBAC4 for 30 minutes in a 37^0^C shaker incubator. The DiBAC_4_ treated bacterial cells were analyzed by flow cytometry (BD FACSVerse by BD Biosciences-US) to evaluate the porosity of the bacterial outer membrane.

### Statistical analysis

Each experiment has been independently repeated 2 to 3 times [as mentioned in the figure legends. The *in vitro* data and the results obtained from cell line experiments were analyzed by unpaired student’s *t*-test by GraphPad Prism 8.4.3 (686) software, and *p* values below 0.05 were considered significant. The results are expressed as mean ± SEM. Differences between experimental groups were deemed to be significant for *p*< 0.05.

## Abbreviations

STM: *Salmonella* Typhimurium
OmpA: Outer membrane protein A
LC3B: Microtubule-associated protein 1A/ 1B-light chain 3
Stx17: Syntaxin 17
LLO: Listeriolysin O
SCV: *Salmonella* containing vacuole
LAMP-1: Lysosome associated membrane protein-1
EEA1: Early endosome antigen 1
RFP: Red fluorescent protein
GFP: Green fluorescent protein
SipC: *Salmonella* invasion protein C

## Author Contributions

ARC and DC conceived the study and designed the experiments. ARC performed all the experiments, analyzed the data, and wrote the original draft of the manuscript. DH performed the experiments, participated in proofreading and editing of the manuscript with ARC. DC supervised the study and reviewed the manuscript. All the authors have read and approved the manuscript.

## Acknowledgments

We sincerely thank the departmental confocal facility, departmental real-time PCR facility, central bioimaging facility, central flow cytometry facility at IISc. Mr. Puneeth and Ms. Navya from the departmental confocal facility are acknowledged for their assistance in image acquisition. Ms. Leepika and Ms. Sharon from the central flowcytometry facility are duly acknowledged for their help in flow cytometry data acquisition. Professor Mohd. Ayub Quadri from the National Institute of Immunology, Newdelhi, is gratefully acknowledged for providing the anti-*Salmonella* SipC antibody. Dr. Shivjee Sah from Professor Umesh Varshney’ lab MCB, IISc is duly thanked for constructing the *ompA*-pQE60 recombinant plasmid.

## Availability of data and materials

All data generated and analyzed during this study, including the supplementary information files, have been incorporated in this article. The data is available from the corresponding author on reasonable request.

## Declarations

### Ethics statement

Not applicable.

### Consent for publication

Not applicable.

## Supplementary Figures

**Figure S1.**
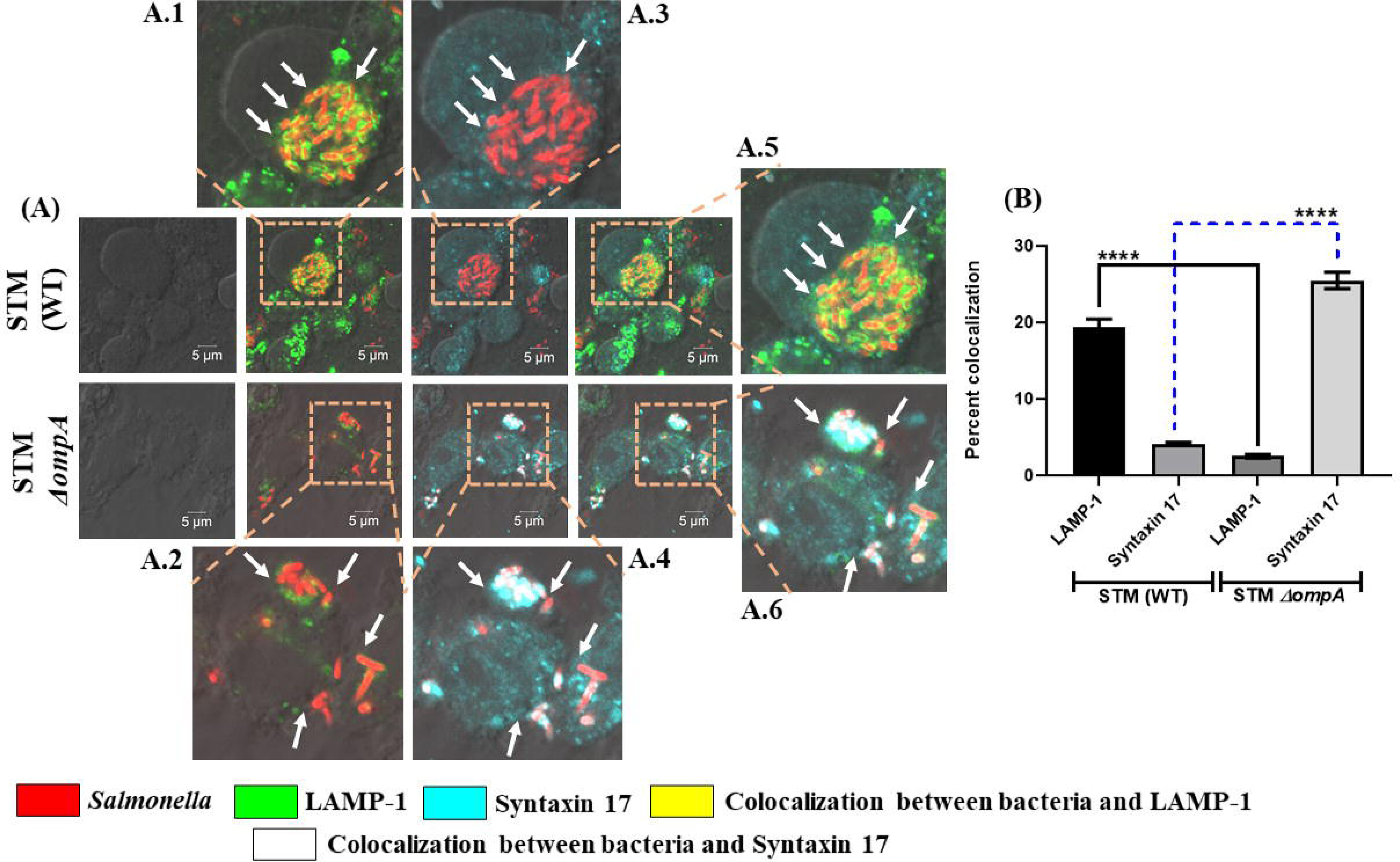
The damage of SCV by STM *ΔompA* activates host autophagy machinery. (A) RAW264.7 cells were infected with STM (WT): RFP and *ΔompA*: RFP at MOI of 20. Cells were fixed at 16 hours post-infection. LAMP-1 and syntaxin 17 were labeled with rat-raised anti-mouse LAMP-1 and rabbit-raised anti-mouse syntaxin 17 primary antibodies, respectively. (B) The quantification of LAMP-1 and syntaxin 17 recruitment on bacteria in RAW 264.7 cells have been represented in the form of a graph. Percent colocalization was determined after analyzing more than 150 different microscopic stacks from two independent experiments. Scale bar = 5μm, [n≥150, N=2]. ***(P)* ****< 0.0001, ns= non-significant, (Student’s *t-*test).**

**Figure S2.**
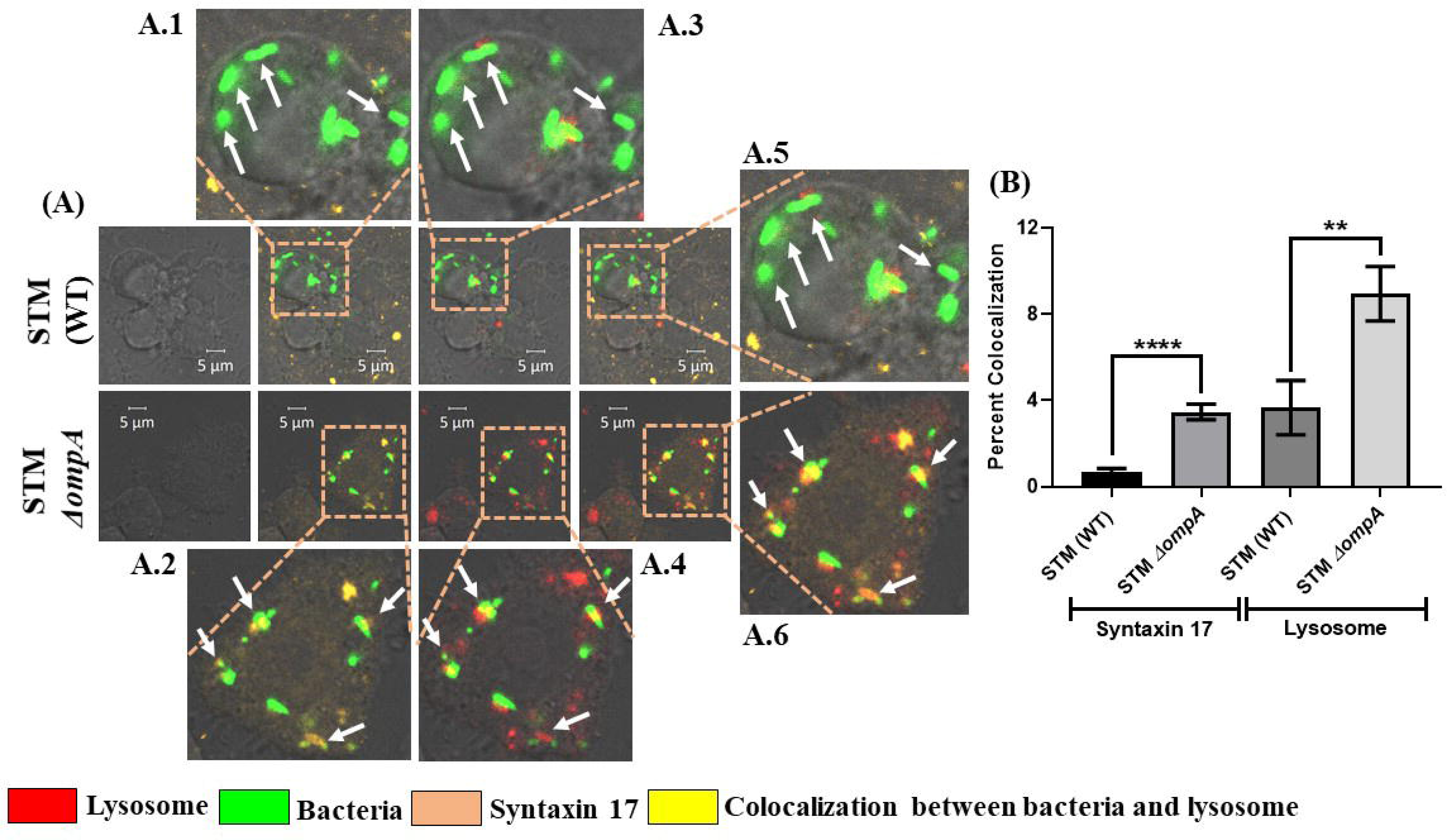
Activation of host autophagy machinery targets the *ompA* deficient *Salmonella* to the lysosome. To stain the lysosomes, RAW 264.7 cells were pre-treated with Texas red ovalbumin for 30 minutes and infected with STM (WT): *GFP* and *ΔompA*: *GFP,* respectively, at MOI of 20. Host syntaxin 17 was labeled with rabbit-raised anti-mouse syntaxin 17 primary and anti-rabbit alexa fluor 647 secondary antibodies, respectively. (H) The colocalization of bacteria with lysosomes and syntaxin 17 has been represented in the form of a graph. The percent colocalization of the bacteria with texas red and syntaxin 17 were determined after counting more than 50 microscopic stacks from two independent experiments (n≥ 50, N=2). Scale bar = 5μm. ***(P)* **< 0.005, *(P)* ****< 0.0001, ns= non-significant, (Student’s *t-*test).**

**Figure S3.**
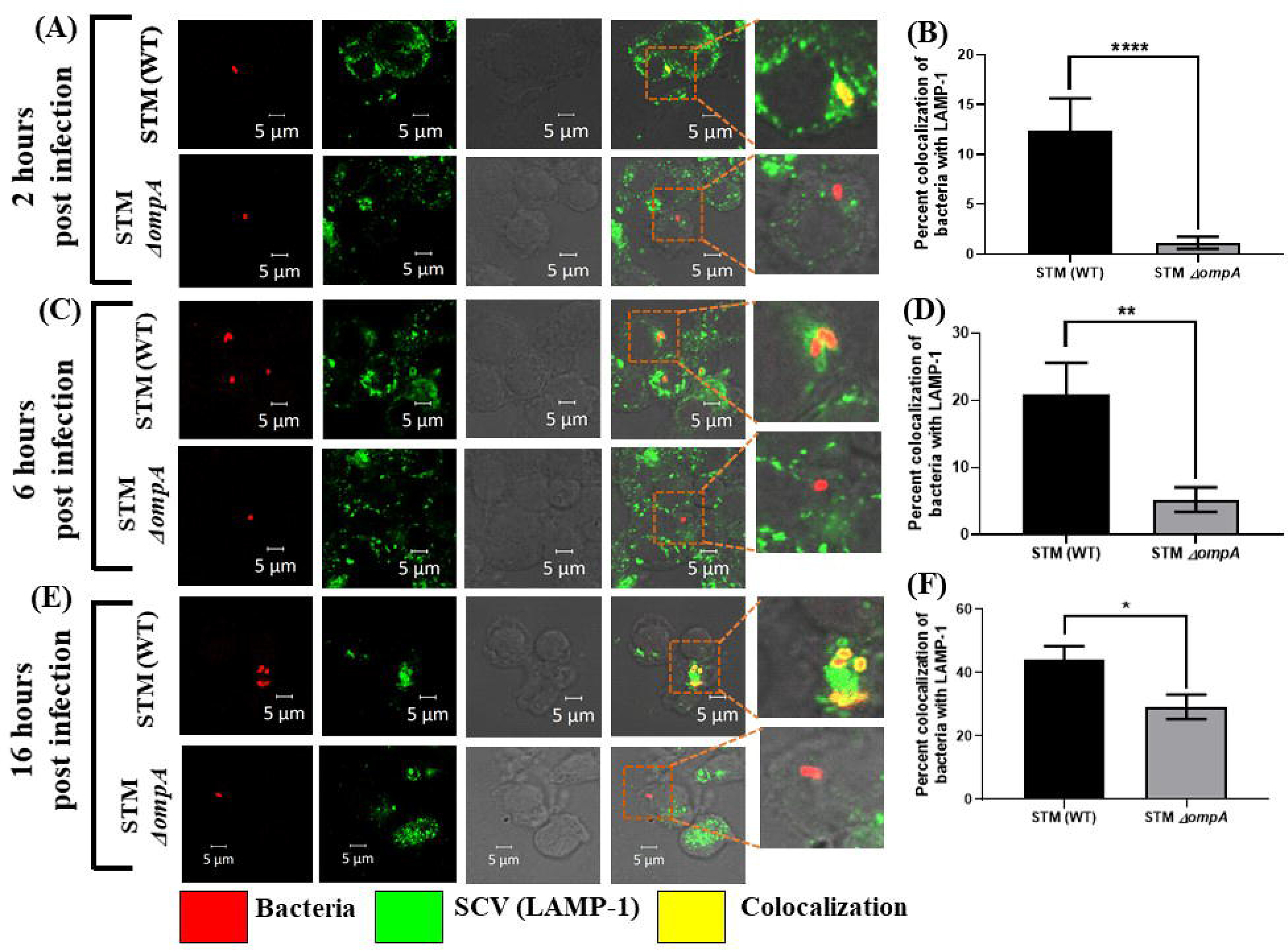
STM *ΔompA* quits the SCV in human monocyte-derived macrophages before the early stage of infection. PMA activates U937 cells were infected with STM (WT): RFP, and *ΔompA*: RFP at MOI of 20. Cells were fixed at (A) 2 hours (early phase), (C) 6 hours (middle phase), and (E) 16 hours (late phase) post-infection & LAMP-1 were labeled with anti-human LAMP-1 antibody. The quantification of LAMP-1 recruitment on bacteria in U937 cells at (B) 2 hours, (D) 6 hours, and (F) 16 hours post-infection has been represented in the form of three graphs. (B) During the early stage of infection (2 hours post-infection), the percent colocalization of bacteria with LAMP-1 was determined after analyzing more than 40 different microscopic stacks from two independent experiments [n≥40, N=2]. (D) During the middle stage of infection (6 hours post-infection), the percent colocalization of bacteria with LAMP-1 was determined after analyzing more than 30 different microscopic stacks from two independent experiments [n≥30, N=2]. (F) During the late stage of infection (16 hours post-infection), the percent colocalization of bacteria with LAMP-1 was determined after analyzing more than 50 different microscopic stacks from two independent experiments [n≥50, N=2]. Scale bar = 5μm. ***(P)* *< 0.05, *(P)* **< 0.005, *(P)* ****< 0.0001, ns= non-significant, (Student’s *t-*test).**

**Figure S4.**
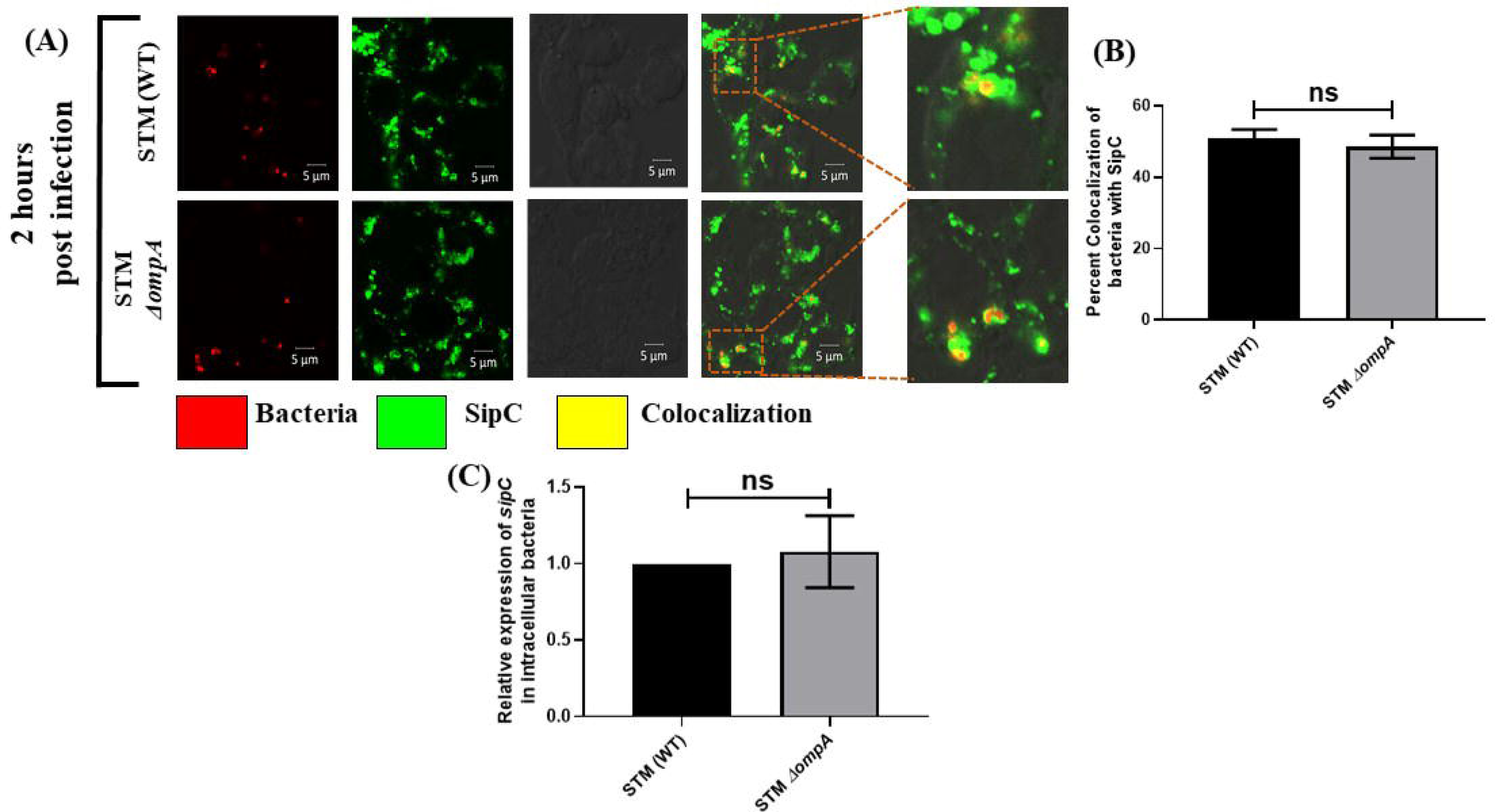
The inability of STM *ΔompA* to retain LAMP-1 does not depend upon the production of SipC. RAW264.7 cells were infected with STM (WT): RFP, and *ΔompA*: RFP at MOI of 20. (A) Cells were fixed at 2 hours post-infection & SPI-1 effector protein SipC produced by intracellular *Salmonella* was labeled with anti-mouse SipC antibody. (B) The quantification of SipC arrangement around the bacteria in RAW264.7 cells at 2 hours post-infection has been represented in the form of a graph. (B) The percent colocalization of bacteria with LAMP-1 was determined after analyzing 100 different microscopic stacks from two independent experiments [n=100, N=2]. Scale bar = 5μm. (C) The transcript level expression of *sipC* in STM (WT) and *ΔompA* growing intracellularly in RAW264.7 cells 12 hours post-infection (n=5, N=3). **ns= non-significant, (Student’s *t-*test).**

**Figure S5.**
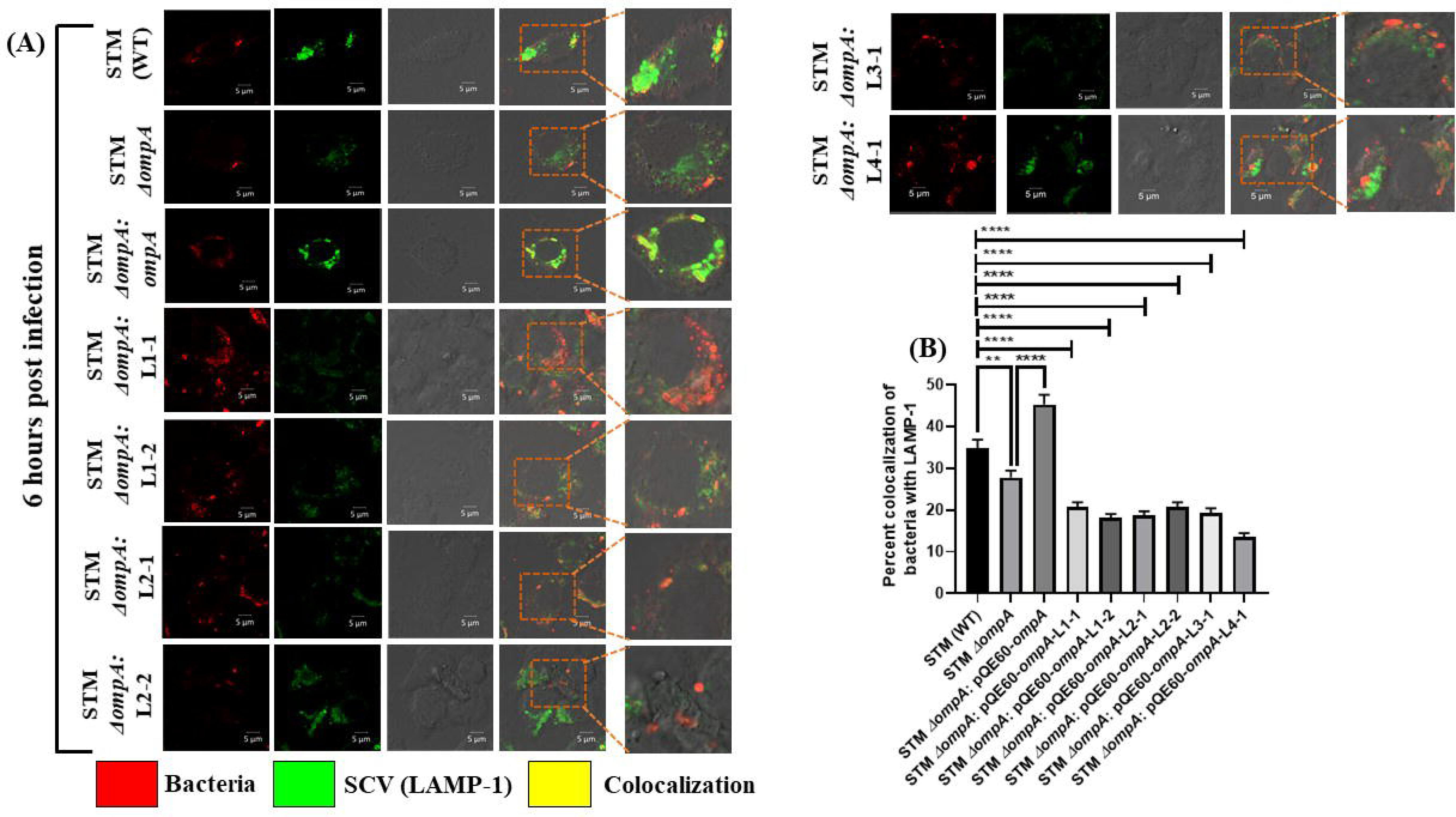
Mutation in the extracellular loops of *Salmonella* Typhimurium OmpA reduces the retention of LAMP-1 around the bacteria in murine macrophages. RAW264.7 cells were infected with STM (WT), *ΔompA*, *ΔompA*: pQE60-*ompA*, *ΔompA*: pQE60-*ompA-*L1-1, *ΔompA*: pQE60-*ompA-*L1-2, *ΔompA*: pQE60-*ompA-*L2-1, *ΔompA*: pQE60-*ompA-*L2-2, *ΔompA*: pQE60-*ompA-*L3-1 and *ΔompA*: pQE60-*ompA-*L4-1 at MOI of 20. Cells were fixed at (A) 6 hours (late phase) post-infection. Intracellular *Salmonella* was stained with rabbit-raised anti-*Salmonella* O primary antibody. LAMP-1 was labeled with rat- raised anti-mouse LAMP-1 primary antibody. The quantification of LAMP-1 recruitment on bacteria in RAW 264.7 cells at 6 hours post-infection has been represented in the form of a graph. (B) During the late stage of infection (6 hours post-infection), the percent colocalization of bacteria with LAMP-1 was determined after analyzing 100 different microscopic fields from two independent experiments [n=100, N=2]. Scale bar = 5μm. ***(P)* **< 0.005, *(P)* ****< 0.0001, ns= non-significant, (Student’s *t-*test).**

